# immgenT: A Comprehensive Reference of Convergent T-cell States in the Mouse

**DOI:** 10.64898/2026.01.30.702892

**Authors:** Ian Magill, Odhran Casey, Dania Mallah, Siba Smarak Panigrahi, Leon Zhou, Olga Barreiro, Derek Bangs, Gavyn Chern Wei Bee, Samantha Borys, Josh Choi, Can Ergen-Mehr, Enxhi Ferraj, Maurizio Fiusco, Antoine Freuchet, Giovanni Galletti, Anna-Maria Globig, Taylor Heim, Charlotte Imianowski, Rocky Lai, Zhitao Liang, Andrea Lebron Figueroa, Erin D. Lucas, Julia Merkenschlager, Kevin Osum, Shannelle P. Reilly, Tomoyo Shinkawa, Claire E. Thefaine, Eric S. Weiss, Liang Yang, Shanshan Zhang, Andre L. Zorzetto-Fernandes, Josh D. Croteau, Maria-Luisa Alegre, Samuel M. Behar, Remy Bosselut, Laurent Brossay, Ken Cadwell, Alexander Chervonsky, Laurent Gapin, Sara E. Hamilton, Jun R. Huh, Iliyan D. Iliev, Bana Jabri, Steve Jameson, Marc K. Jenkins, Susan M. Kaech, Daniel H. Kaplan, Vijay Kuchroo, Michel Nussenzweig, Marion Pepper, David A. Sinclair, Stefani Spranger, Michael Starnbach, Matthew Stephens, Ulrich von Andrian, Nir Yosef, Joonsoo Kang, Mitch Kronenberg, Ananda Goldrath, Maria Brbić, Christophe Benoist, David Zemmour, the immgenT Project

**Author notes:** **Address correspondence to:** David Zemmour, MD, PhD 927 East 57th St, Chicago, IL 60637.

## Abstract

The immgenT collaborative project generated a comprehensive molecular atlas of T cells spanning virtually all mouse organs and disease states, profiling ∼800,000 cells from 750 samples with RNA, 128-plex surface protein, and αβTCR sequence. Applying a deep generative model to joint RNA and protein data defined the landscape of T-cell states organized into eight lineages and 107 robust clusters, integrating similar cells from different contexts, and resolving prior nomenclatures. Analysis of effector molecules, transcription factors and modules showed that both immunological functions and regulatory programs are shared across cell states. This framework provides a stable, reusable reference, demonstrated by computationally integrating 16 external datasets from diverse biological contexts. A set of public web tools supports browsing of these data and mapping of any dataset onto the immgenT framework. These results propose a molecular classification of T cells organized around a set of shared states reused across immunological contexts.

## Introduction

Our understanding of T lymphocytes has a long and rich history^1^. From humble beginnings as products of a vestigial organ with no discernible function in immunity, T cells became recognized as indispensable helpers of antibody responses^2^, and mediators of anti-viral defense, with the bewildering notion of “restriction” by molecules of the Major Histocompatibility Complex (MHC)^3,4^, a perplexity resolved by the demonstration of presentation of peptidic antigens in the groove of MHC molecules^5^.

T cell’s recognized influence has grown to encompass essentially all immunological modalities, with the growing perception that T cells patrol and help maintain the homeostasis of many non-immunological organ in the mammalian body. Accompanying these diverse functions, the field has partitioned T cells into specialized subsets, starting with the fundamental division into CD4^+^ and CD8^+^ lineages^6^ restricted by MHC-I vs MHC-II molecules^7^, respectively, followed by splits based on cytokine secretion^8^, the recognition of non-conventional T cell types^9–12^, and ultimately into ever-finer functional states.

T cell diversity has long been ascertained by flow cytometry^13,14^, the original single-cell technology, that also enables functional analysis, with increasingly complex marker panels. Although still the gold standard, flow cytometry approaches have run into trouble: even with finely sorted pools of cells based on a 20+ marker panels, insufficiently resolved populations can confound functional inferences. Immunologists are decried as splitters by other biologists, forever subsetting their populations with thin justification. Conversely, conclusions of functional correlates based on too few markers, and their inferred relationships in variable contexts, have led to confusion. In the last decade, single-cell transcriptomics has opened the door to parsing T cells based on thousands of gene products, with the explicit promise of fully resolving the scope of T cell states^15^. Unfortunately, single-cell transcriptomics may have made matters worse. Because each experiment is limited to a selected organ, challenge condition or cell-type, and because outputs of single-cell computational tools do not transport readily, each experiment tends to live in its own universe. Every study generates its own cartography (the typical “UMAP”) that is never reused elsewhere, with ad-hoc annotations that give up on a unified understanding. Some studies have made concerted efforts to provide a unified framework for some lineages^16–19^, and there are computational tools that could be applied to make sense of the diverse accumulated data^20–25^. However, there is no coherent cosmology into which a T cell study can be inscribed, and instead a proliferation of subsets and acronyms that have variable support and reliability, as illustrated by a recent perspective^26^.

The 41 labs who participated in the “immgenT” Open Source Program^27^, in line with the ImmGen consortium’s practice of applying common operating protocols^28^ to generate broad community references, set out to coordinately profile T cells across all body locations in the mouse and across the spectrum of T cell states induced by immunologic challenge. Single-cell RNAseq, CITEseq^29,30^ and TCRseq were used coordinately. A broadly and rigorously generated dataset of this scale should reveal the extreme states that T cells can adopt, and the regulatory programs that underlie them. It should allow the identification of unrecognized states, clarify redundancies and commonalities in cell phenotypes and transcriptional programs, and identify actionable combinations of cell-surface markers. Perhaps most importantly, this effort could yield a useful framework of T cell states against which other T cell studies could be compared.

The present article presents a general overview of the results obtained by the immgenT program, in relation to these goals. It is accompanied by focused satellite articles that distill these learnings about specific lineages and conditions. Together, these data define—through AI-based joint RNA–protein embedding—a structured molecular landscape of mouse T cells, comprising eight major lineages and 107 reproducible states that recur across tissues and immune perturbations. Beyond defining this landscape, we introduce a practical online tool that enables direct comparison across studies. By anchoring a reusable reference embedding and consensus annotations, immgenT allows new datasets to be integrated without recomputing dimensionality reductions. In addition the CITEseq data provides a direct bridge between high-dimensional molecular states and routine flow cytometry–based immunology. We further demonstrate that this framework generalizes to external datasets and supports cross-study harmonization. In this sense, immgenT introduces a new T-cell cosmology: a coherent, extensible map into which diverse T-cells, past and future, can be placed, compared, and interpreted within a unified molecular framework.

## Results

### immgenT

The immgenT OpenSource Project is a community-driven effort to build a comprehensive molecular profile of T cells across tissues and immune perturbations (**Extended Data Table 1**). The project brought together more than 100 investigators across over 40 laboratories, with experiments coordinated through shared standard operating procedures and centralized data processing. All data and analytical resources are released openly, in keeping with the ImmGen tradition of reproducibility and community access (see Methods for description of the web portals).

The consortium performed single-cell RNAseq with a standardized CITEseq framework^29,30^ to jointly measure mRNA and surface protein expression, leveraging a collaboratively developed 128-antibody pconsanel (BioLegend) (**Extended Data Table 2**), with paired αβ TCR sequencing (**Fig. 1a**).

**Figure 1.**
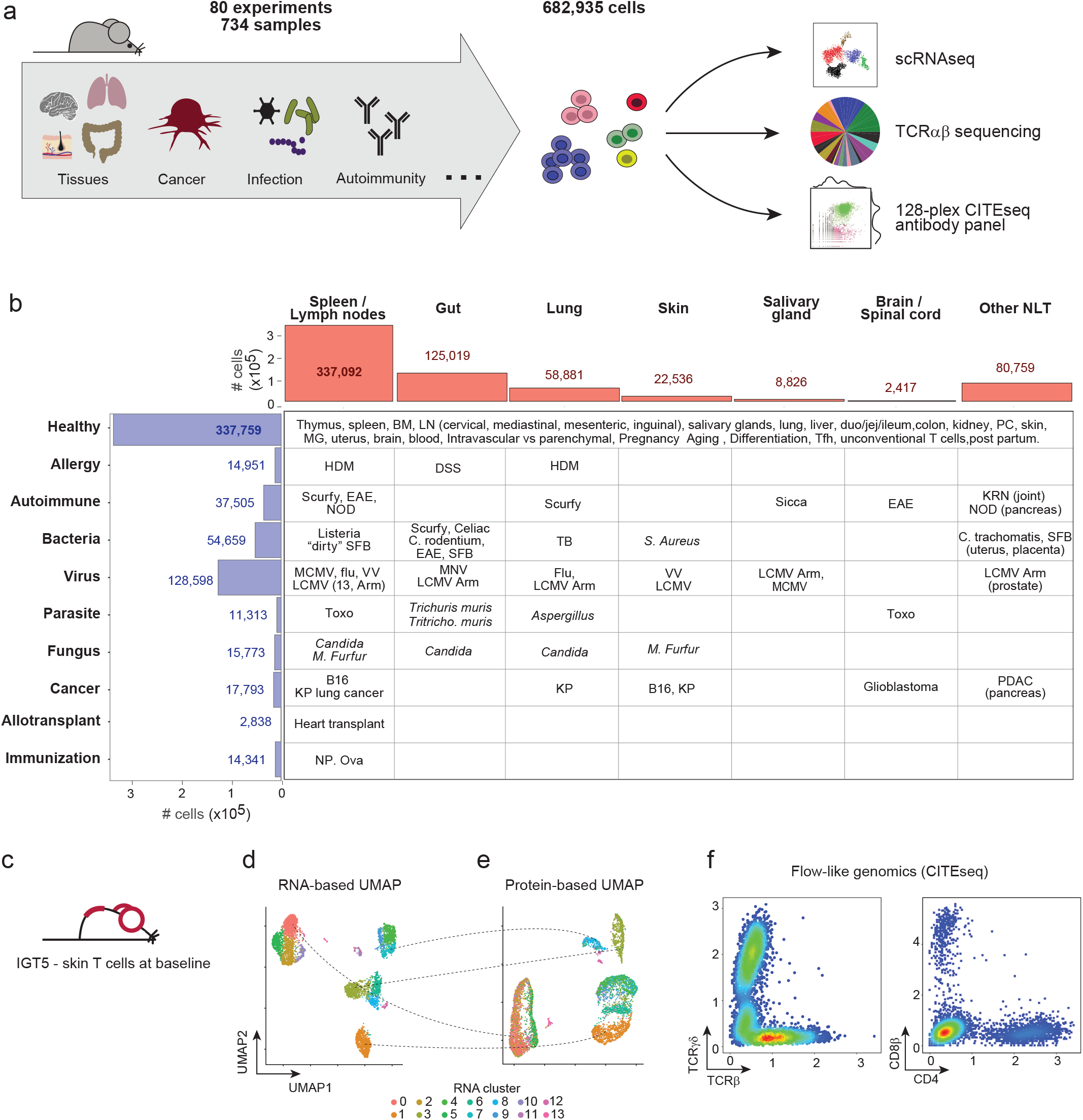
Scope of the immgenT Open Source Project. **(a)** Schematic overview of the immgenT project. Across 80 successful experiments encompassing 734 samples spanning diverse tissues and immune perturbations, a total of 682,935 T cells were profiled using single-cell RNA sequencing, paired αβ TCR sequencing, and CITE-seq with a 128-antibody surface protein panel. **(b)** Summary of sampled conditions organized by tissue and broad disease category. Bar plots indicate the total number of T cells sequenced per organ (top) and per disease category (left). **(c–f)** Representative immgenT dataset (IGT5), profiling skin (ear and back) at baseline conditions. **(d)** UMAP visualization based on gene expression, with cells colored by transcriptomic clusters. **(e)** UMAP based on surface protein expression (CITE-seq), with cells colored by the gene-expression clusters shown in (d). **(f)** Flow-cytometry–like expression of TCRγδ versus TCRβ and CD4 versus CD8β expression (CITEseq, log1p(CP10K)). **Abbreviations**: BM, bone marrow; CNS, central nervous system (brain and spinal cord); LN, lymph node; MG, mammary gland; NLT, non-lymphoid tissue; PC, peritoneal cavity; PLO, primary lymphoid organ; SLO, secondary lymphoid organs. Disease models: B16 (B16 melanoma), *C. trachomatis* (*Chlamydia trachomatis*), DSS (dextran sodium sulfate colitis), EAE (experimental autoimmune encephalomyelitis), HDM (house dust mite), KP (Kras-p53–driven cancer), KRN (K/BxN arthritis), LCMV (lymphocytic choriomeningitis virus, Armstrong and Clone 13), MCMV (murine cytomegalovirus), MNV (murine norovirus), NOD (non-obese diabetic), NP-OVA (NP-ovalbumin immunization), PDAC (pancreatic ductal adenocarcinoma), *S. aureus* (*Staphylococcus aureus*), SFB (segmented filamentous bacteria), TB (*Mycobacterium tuberculosis*), Tritrich. muris *(Trichichomonas muris)*, Toxo (*Toxoplasma gondii*), and VV (vaccinia virus).

The dataset is almost exclusively based on C57BL/6 mice and spans more than 46 distinct tissues and anatomical sites (**Fig. 1b**). Samples were collected both at baseline (6–8-week-old male or female specific-pathogen–free mice) and across diverse physiological and immunological conditions, including aging, pregnancy and postpartum states, immunization, autoimmunity and inflammation (e.g., NOD insulitis, DSS colitis, Foxp3-deficiency), tumors, and a range of acute and chronic infections (*Chlamydia trachomatis* in the uterus, *Toxoplasma gondii* in the brain, and *Mycobacterium tuberculosis* in the lung).

T cells were defined operationally as cells expressing a TCR and were therefore identified primarily by CD3 expression. Most experiments sequenced total CD3⁺ T cells to capture both well-characterized and potentially unanticipated populations. To ensure adequate representation of rare subsets, targeted experiments were also performed, including enrichment for CD4+ FoxP3⁺ regulatory T cells, unconventional T cells (e.g., Zbtb16^+^ cells, or CD4⁺ T cells in MHC class II–deficient mice and CD8⁺ T cells in MHC class I–deficient mice), and CXCR5^hi^ PD-1^hi^ T cells following ovalbumin (OVA) immunization. Antigen-specific T cells were profiled whenever feasible, particularly in viral, bacterial, and tumor models, using tetramer staining (either sorted or captured via DNA-barcoded CITEseq reagents) or adoptive transfer of TCR-transgenic cells (e.g., P14 in the B16-GP33 tumor model). Experiments did not encompass natural or experimental genetic variation, except when necessary to anchor disease-relevant populations (PD-1–deficient T cells^31^, *Foxp3*-deficiency (*scurfy*), NOD diabetes).

Altogether, the project comprises 80 datasets (referred as “IGT”), each designed as a standalone study with appropriate internal controls (**Extended Data Table 1**). Within each experiment, up to 20 samples were hashtagged and pooled prior to encapsulation, enabling efficient recovery of rare populations while limiting batch effects. Each experiment included a standard reference sample (splenocytes from a 6–8-week-old male or female C57BL/6 mouse from the Jackson Laboratory) to enable cross-experiment normalization and assessment of technical variability. After quality control (**Extended Data Note 1, Extended Data Fig. 1, Extended Data Fig. 2, Extended Data Table 3, Extended Data Table 4**), the dataset includes 682,951 cells across 734 samples, including 42,114 antigen-specific T cells. Forty different organs were sampled, spleen and lymph nodes most extensively, followed by gut and lung (**Fig. 1b**), with balanced female/male representation. The protein data were of good quality for 102 of 128 markers attempted (**Extended Data Note 1)**, with notable failures for some chemokine receptors, which require different staining conditions.

To illustrate the data, we highlight IGT5, profiling of ear and back skin at steady-state (**Fig. 1c-f**). Cells formed well-resolved clusters in RNA-based Uniform Manifold Approximation and Projection (UMAP) space^32^ (**Fig. 1d**), and these same clusters remained clearly separable when cells were embedded using only surface protein expression measured by the 128-antibody CITEseq panel (**Fig. 1e**). This concordance illustrates that the surface markers included in the panel capture sufficient information to preserve transcriptionally defined T cell states (**Extended Data Note 1**). In addition, protein-level data enabled flow cytometry–style analyses, as illustrated by the scatter plots of **Fig. 1f**, which closely resemble flow cytometry results.

Together, these analyses illustrate how each immgenT experiment functions as a self-contained dataset that can be explored through both high-dimensional single-cell analysis and traditional immunological frameworks. To facilitate broad access and interactive exploration, we developed Rosetta2, a web-based interface that enables users to browse, visualize, and analyze each immgenT experiment independently (**Extended Data Note 2**).

### immgenT harmonizes T cell representation across experiments and delineates major T cell lineages

Moving beyond individual experiments to enable meta-analysis required integrating data from all 80 datasets and 734 samples into a shared representation. To this end, we applied totalVI^25^, an deep generative AI model for data integration that jointly utilizes RNA and surface protein abundance to learn a shared latent representation^20^. To retain maximal biological information, we used the full set of measured genes and CITEseq markers (re-running the integration with a subset of variable genes yielded highly similar results). The integrated latent space was subsequently projected using minimum distortion embedding^33^ (MDE) to a 2D-representation of all T cells that incorporates both their RNA and protein characteristics (hereafter “all-T MDE”) (**Fig. 2a**). We used MDE instead of the more common UMAP to allow new data to be integrated while maintaining the same embedding.

**Figure 2.**
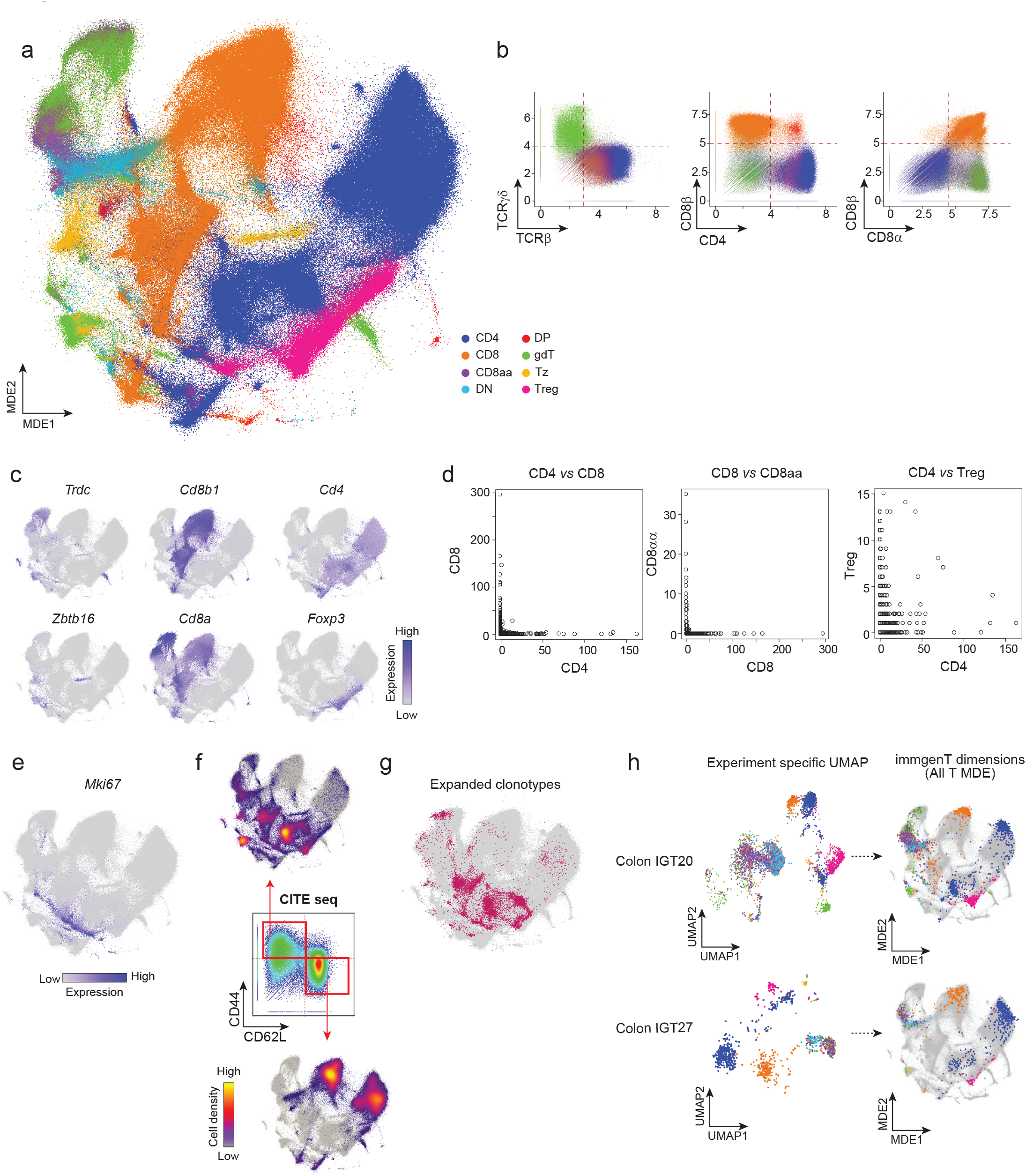
immgenT harmonizes T-cell representation across experiments and delineates major T-cell lineages. **(a)** Reference MDE plot of all profiled T cells (all-T MDE), colored by T-cell lineage. CD4, conventional CD4⁺ T cells; CD8, conventional CD8αβ⁺ T cells; Treg, CD4⁺ Foxp3⁺ regulatory T cells; CD8aa, CD8αα αβ T cells; gdT, γδ T cells; Tz, Zbtb16⁺ αβ T cells; DN, CD4⁻CD8αβ⁻ αβ T cells; DP, CD4⁺CD8αβ⁺ αβ T cells. **(b)** Expression of cell surface markers in the CITEseq data (log1p(CP10K)). Cells are colored by lineage as in (a). **(c)** all-T MDE color coded by RNAexpression of canonical lineage transcripts (*Trdc, Cd8b1, Cd4, Zbtb16, Cd8a,* and *Foxp3*). **(d)** TCRαβ clonotype sharing between CD4, CD8, CD8a, and Treg populations (number of occurences across the entire non-transgenic dataset) **(e)** *Mki67* expression in the all-T MDE, highlighting proliferating cells. **(f)** All-T MDE with CD62L⁺CD44⁻ and CD44⁺CD62L⁻ T cells highlighted, gated as shown in the adjacent CD62L versus CD44 protein expression plot (CITEseq, log1p(CP10K)). **(g)** all-T MDE highlighting expanded clonotypes identified by αβ TCRseq **(h)** Colonic T cells at baseline profiled in two independent experiments (IGT20, top; IGT27, bottom). Left, pre-integration UMAPs computed separately for each dataset. Right, post-integration representation of the same cells in the all-T MDE, colored by lineage as in (a).

The design of immgenT included an internal standard (normal splenocytes) spiked into each experiment. Each of these overlapped tightly in the integrated space (**Extended Data Fig. 3a**), indicating minimal residual batch effects following integration.

Across all tissues and perturbations, the all-T MDE resolves major T cell lineages as large, contiguous regions, or “continents” (**Fig. 2a, Extended Data Fig. 3b**), rather than segregating primarily by tissue of origin or experimental condition. Based on the expression of specific protein (**Fig. 2b**) or RNA markers (**Fig. 2c**), T cells were parsed into lineages (“level1” annotation). Conventional CD4⁺ T cells (“CD4”), conventional CD8αβ⁺ T cells (“CD8”), and CD4⁺ FOXP3⁺ regulatory T cells (“Treg”) formed clearly separated regions in the embedding (**Fig. 2a, Extended Data Fig. 3b**). Annotation using CITEseq surface markers (anti-TCRγδ,-TCRβ,-CD4,-CD8α, and-CD8β) further revealed that 23% of cells in the dataset (148,286 cells) fell outside of these populations and comprised γδ T cells (“γδT”), CD8αα⁺ TCRαβ⁺ T cells (“CD8aa”), as well as mature CD4⁻CD8⁻ double-negative (“DN”) and CD4⁺CD8⁺ double-positive (“DP”) populations (**Fig. 2b,c, Extended Data Fig. 3c**). Lastly, a sixth population of Zbtb16⁺ TCRαβ⁺ T cells (“Tz”) formed distinct regions with variable expression of CD4 and CD8 (**Fig. 2a, Extended Data Fig. 3b,c**). Paired αβTCR sequencing identified the canonical TCRs of invariant NKT^34^ (6,214 cells) and of MAIT cells^12^ (438 cells) (**Extended Data Fig. 3d,e**), all mapping to the Tz population (further described in the immgenT-Tz paper).

These distinctions into high-level lineages reflect fundamental differences in thymic differentiation pathways, driven by distinct MHC restriction and TCR signaling requirements during differentiation^35^. Consistent with this interpretation, analysis of the TCRαβ repertoire showed essentially no overlap between CD4 and CD8 lineages, whereas CD4 and Treg lineages showed greater clonotype overlap (**Fig. 2d**), consistent with earlier results^36,37^, and reflecting their shared restriction by MHC-II molecules and known interconversion (peripherally induced Tregs). Beyond lineages, other factors seem to organize the distribution of the cells in the latent space. In particular, proliferating cells clustered separately from the non-proliferating cells in each lineage, as shown with *Mki67* expression (**Fig. 2e, Extended Data Fig. 3f**). Resting and activated states highlighted with CD62L and CD44 expression also clearly separate cells across lineages (**Fig. 2f**). As might have been expected, cells that have undergone clonal expansion, as shown by their shared TCR clonotypes, are found in the activated clusters (**Fig. 2g**).

An intriguing subset of cells (5.7%) consistently clustered into a distinct state, which we termed “miniverse” (wM clusters) (**Extended Data Fig. 3f**). Miniverse cells included representatives from all lineages and experimental conditions and were characterized by a low-translation, low-mitochondrial, G2/M-arrested program, suggestive of stressed cells arising from an unresolved biological state, or a technical artifact. Such populations are commonly observed in single-cell RNAseq datasets and were retained here to preserve the completeness of the immgenT reference atlas.

This integration replaces dozens of experiment-specific embeddings with a unified reference space. For example, two datasets of colon T cells (IGT20, 27) analyzed independently yield UMAP representations difficult to relate (**Fig. 2h, left**). Following integration, cells from both experiments overlap in the all-T MDE, with CD4, CD8, γδ T, and other T cell lineages aligning consistently across datasets (**Fig. 2h, right**). Together, these results establish a unified, lineage-centered reference space for T cells, in which intrinsic developmental programs dominate over tissue and condition-specific variation, providing a foundation for comparative analysis of T cell states across diverse immune contexts.

### Integration into 107 consensus clusters preserves heterogeneity and enhances biological interpretability

While the integrated all-T MDE provides a global reference space for distinguishing major lineages, there was also finer-grained heterogeneity within individual lineages. Re-clustering in the integrated latent space yielded 107 clusters, representing a consensus across all datasets. Clustering resolution was optimized using silhouette score analysis^38^ to balance under-and over-clustering with regard to gene signature, sample distribution, surface marker expression (**Extended Data Note 3, Extended Data Fig. 4**). These clusters are more readily visualized on major lineage-specific MDE projections, leading to the representations of **Fig. 3a**. These lineage-level MDE maps have become the operating standard in the immgenT program for fine parsing within lineages, used in this and companion manuscripts, which explore in depth the transcriptional, protein, and contextual features of individual lineages and states. By deference to the Human Leukocyte Antigens (HLA) and cluster of differentiation (CD) traditions, small clusters of unclear significance are given a “w” prefix. Details on the 107 clusters, their organismal distribution and differential gene expression are available at https://www.immgen.org/ImmGenT/ (**Extended Data Note 2**).

**Figure 3.**
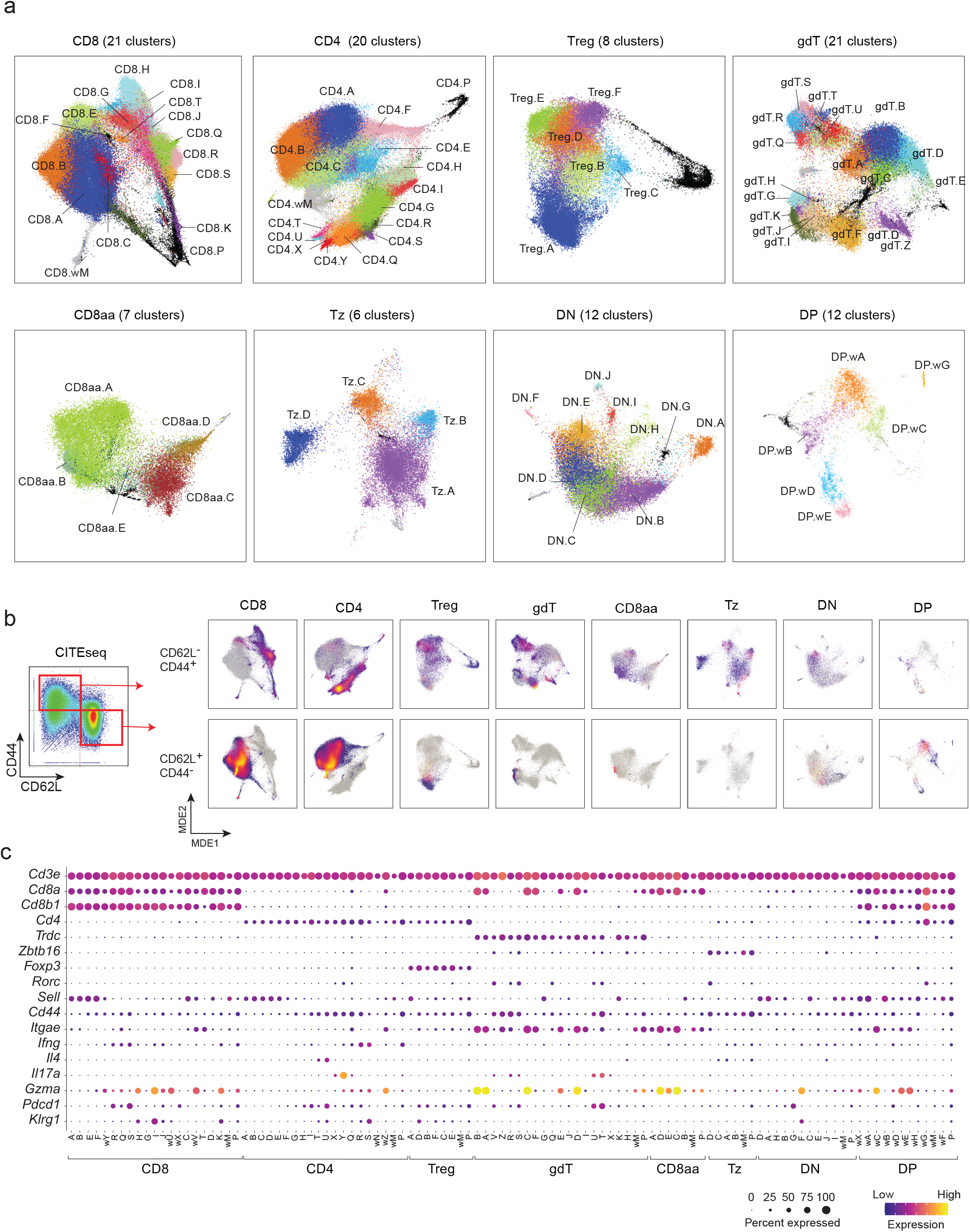
Reference MDE plot for each T-cell lineage. **(a)** Reference MDE plot for each lineage. Colors by clusters. **(b)** CD44⁺CD62L⁻ (top) and CD62L⁺CD44⁻ (bottom) T cells highlighted in each lineage-specific MDE. Gated cells as shown in the adjacent CD62L versus CD44 protein expression plot (CITEseq, log1p(CP10K)). **(c)** Dot plot showing scaled expression of canonical T-cell transcripts across clusters.

These lineage-restricted embeddings recover well-established patterns. For example, expression of the classical markers CD62L and CD44 segregates resting and activated states within CD4⁺ T cells, regulatory T cells, and CD8⁺ T cells, with clusters aligning along these axes (**Fig. 3b**). There was greater heterogeneity within the CD44+ activated and memory states. The expression of genes commonly used for cell identification also recapitulated familiar patterns (**Fig. 3c)**. We also mapped the expression of broad stress-interferon-or hypoxia-related signatures, which are frequently encountered in scRNAseq datasets, and can denote biological variables or technical stress^39–42^. Most showed a broad distribution across the T cell space, and across organs and conditions (**Extended Data Fig. 5a-j**). A striking exception was the Interferon signature, which was dominant in specific states (e.g. CD8.F or CD4.D (**Extended Data Fig. 5f,k)**, in keeping with several reports of small T cell populations with high ISG expression^43,44^.

Several strategies were employed to address whether the 107 consensus clusters (where “consensus” reflects that they originate from many independent samples and conditions) provide a faithful and sufficiently granular representation of T-cell heterogeneity or whether integration obscures biologically meaningful variation. First, cluster discovery rapidly saturated: all 107 clusters were detected after approximately two-thirds of the experiments had been incorporated (**Fig. 4a**), indicating that the reference captured a near-complete repertoire of states present across the dataset.

**Figure 4.**
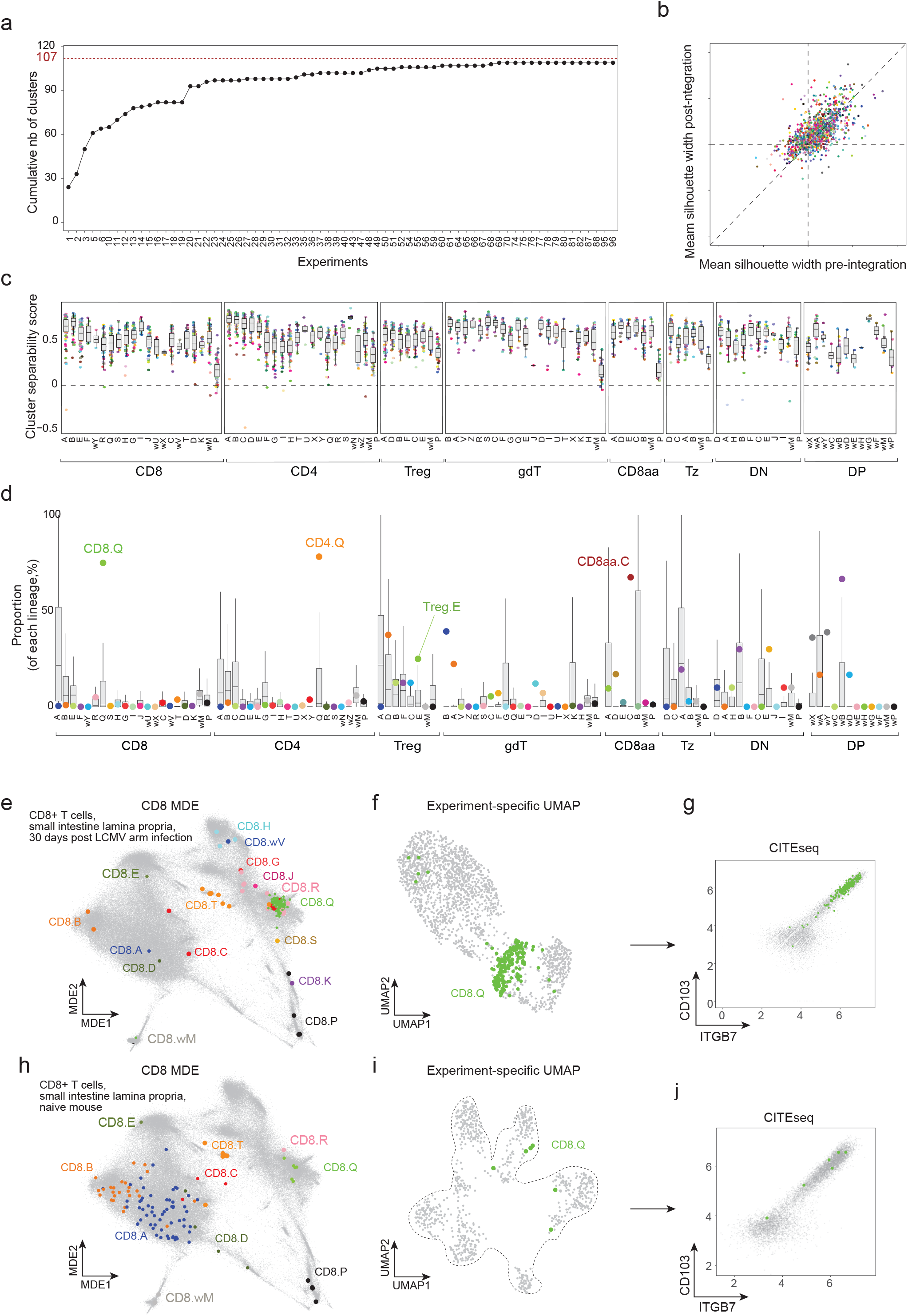
107 immgenT consensus clusters preserve heterogeneity and enhance biological interpretability. **(a)** Saturation of T-cell states within the immgenT dataset. The number of clusters containing at least 50 cells across experiments reaches saturation after the 69th experiment. **(b)** Cluster separation (mean silhouette width) before integration (within each dataset) and after integration into the immgenT latent space, showing the extent to which well-separated clusters remain distinct following integration. Colors indicate experiment (IGT). Scores > 0 denote well-separated clusters. R² = 0.38 **(c)** Distribution of cluster separability scores (see Methods) for each immgenT consensus cluster across experiments in the gene expression space. Scores > 0 indicate transcriptionally distinct clusters. Colors indicate experiment (IGT). **(d–j)** Contextualizing individual experiments within the immgenT reference enhances biological interpretability. **(d–g)** T cells from the small intestine 30 days after LCMV Armstrong infection, colored by immgenT clusters. **(d)** Proportion of each immgenT cluster within each lineage in the sample. Background boxplots show the distribution of cluster proportions across all samples, highlighting clusters enriched in this sample (most CD8 T cells map to CD8.Q). **(e)** Sample’s CD8 T cells in the CD8-specific MDE, with all immgenT CD8 T cells shown in grey for reference. **(f)** Experiment-specific UMAP, showing that cells assigned to immgenT CD8.Q in the sample cluster together. **(g)** Sample’s CD8.Q cells express tissue-resident memory markers CD103 and ITGB7 (CITEseq, log1p(CP10K)). All immgenT CD8 T cells are shown in grey. **(h–j)** T cells from the small intestine of a naïve mouse, colored by immgenT clusters. **(h)** Sample’s CD8 T cells in the CD8-specific MDE, with all immgenT CD8 T cells shown in grey. CD8.Q cells are rare at baseline. **(i)** Experiment-specific UMAP, where the rare CD8.Q cells do not form a distinct cluster. **(j)** Sample’s CD8.Q cells express CD103 and ITGB7 (CITEseq, log1p(CP10K)). All immgenT CD8 T cells are shown in grey.

Second, integration did not collapse the experiment-specific structures. To assess this, we evaluated whether clusters identified independently within each experiment retained coherent organization in the integrated latent space. Silhouette widths of pre-integration clusters were significantly correlated before and after integration (R² = 0.38; Spearman ρ = 0.63; **Fig. 4b**), demonstrating that clusters that were well defined within individual datasets remained well separated following integration.

Third, we assessed the robustness of consensus clusters independently within each dataset’s transcriptomic and proteomic spaces (**Extended Data Note 3, Fig. 4c, Extended Data Fig. 4c,d**). Notably, 99.5% of clusters showed positive separability in RNA and 95% in protein, supporting their reproducibility and molecular distinctness.

Finally, rather than masking biological differences, mapping cells to the immgenT reference augmented each dataset and enabled two complementary analyses: (i) identification of T-cell clusters that are selectively enriched under specific conditions and therefore likely to be biologically relevant, and (ii) detection of rare T-cell states that are difficult to resolve in standalone analyses. A defining feature of the integrated dataset is the distribution of cluster abundances across samples. Although the 107 consensus clusters are shared across many experiments, their relative proportions vary widely. Most clusters are rare or absent in the majority of samples, whereas a small number of samples exhibit strong enrichment—sometimes exceeding 75% of all T cells for a given cluster (**Fig. 4d**). This long-tailed distribution provides a background against which condition--specific expansions can be readily identified.

For example, comparison of small-intestine lamina propria (SI LP) T cells from mice 30 days after LCMV Armstrong infection revealed marked enrichment of specific clusters across multiple lineages, including CD8.Q, CD4.Q, and Treg.E (**Fig. 4d**). CD8.Q was particularly expanded at this memory time point and formed a tight, well-defined cluster in the CD8 MDE (**Fig. 4e**), as well as the original experiment UMAP (**Fig. 4f**). CITEseq analysis showed expression of CD103 and ITGB7, consistent with a tissue-resident memory phenotype (**Fig. 4g**).

In contrast, CD8.Q cells were very rare in a naive mouse SI LP, with as few as six cells detected (**Fig. 4h-j**). Such sparse populations are insufficient to form distinct clusters in standalone analyses and can be easily overlooked or misannotated (**Fig. 4i**). Mapping to immgenT nonetheless assigns these cells to the appropriate consensus cluster based on shared transcriptomic and protein features (**Fig. 4i**), a classification further supported by consistent CD103 and ITGB7 expression (**Fig. 4j**).

Together, these results demonstrate that the 107 immgenT consensus clusters preserve the original complexity of the 734 samples while providing a unified reference framework. By defining a quantitative background distribution of cluster abundances, immgenT facilitates identification of condition-enriched T-cell states and enables reliable detection and annotation of rare populations, extending the interpretability of individual experiments beyond their intrinsic sampling depth.

While a detailed analysis of lineage-specific heterogeneity is presented in companion manuscripts (**immgenT-CD8**^45^**,-Treg**^46^**,-CD4**^47^**,-Tz**), we provide a concise overview here. Within CD4⁺ T cells (see companion immgenT-CD4 manuscript), we observed a model of graded polarization in which fully polarized cytokine-producing clusters (including IFNγ (CD4.R/S), IL-4 (CD4.H/T), and IL-17 (CD4.X/Y)), co-exist with a majority of CD44⁺ cells that occupy a larger population with lower and overlapping expression of these effector cytokines. An additional set of states spanned multiple transcriptional programs and revealed unexpected proximity between Tfh-like CXCR5⁺PD-1⁺ cells arising during immunization (CD4.H) and cells associated with chronic infection or cancer (CD4.I). For CD8⁺ T cells (see **immgenT-CD8 manuscript**^45^), heterogeneity could loosely be matched to labels in common use, with clusters somewhat corresponding to effector (CD8.H/I/J/wU), central memory (CD8.E/F), effector memory (CD8.G), tissue-resident memory (CD8.Q), and exhausted-like states (CD8.R/Q/S). However, these molecular states spanned a wider range of experimental conditions than their conventional labels suggest and cells enriched in chronic infection and cancer showed substantial transcriptional overlap with clusters typically associated with tissue residency^48^. These patterns, explored in greater depth in the accompanying manuscript (**immgenT-CD8**^45^), argue for defining CD8⁺ T-cell states primarily by shared molecular programs rather than by condition-specific labels. Treg cells (see **immgenT-Treg manuscript**^46^) exhibited a more limited heterogeneity, comprising six principal clusters displayed distinct tissue-and condition-specific distributions, including a prominent population of CD49b⁺ Tregs^49^ enriched in blood and the placenta. γδT cells were organized primarily along a major axis separating CD8αα⁺ populations enriched in the gut from CD49A⁺ populations. Unconventional αβ T cells, including MAIT and iNKT cells, overlapped transcriptionally with each other and with other non-conventional T cells across four main states, largely engaging gene programs shared with CD4⁺ T cells (see **immgenT-Tz manuscript**).

### T-RBI integrates external datasets into the immgenT framework

A central goal of immgenT was to establish a robust and comprehensive framework capturing the breadth of T cell heterogeneity. The fact that the 734 immgenT samples could all be integrated into the unified reference space suggested that the model is very general, but assessing the coverage and generalizability of this framework required systematic validation using independent external datasets. To this end, we developed T-RBI (T cell Reference-Based Integration), a reference-mapping strategy that integrates external datasets with the immgenT reference using successive rounds of scVI and scANVI^20,50^. T-RBI produces two complementary outputs: (i) immgenT cluster annotations with associated confidence scores for each query cell and (ii) coordinates of query cells within the fixed immgenT MDE graphs^51^. The public implementation of T-RBI is available at https://rstats.immgen.org/immgenT_integration_analysis/).

To systematically test whether external datasets contain T cell states absent from immgenT, we applied T-RBI to 16 independent studies^52–55^ comprising 335,042 cells (**Extended Data Table 5**). These datasets spanned a wide range of biological contexts, including prior atlas efforts (Tabula Muris^56^ and a CD8 T cell atlas^16^), conditions and settings not included in the immgenT compendium, such as CAR T cells^57^, Tregs in the meninges^58^ and skeletal muscle^59^, AAV–HBV models^60^, and multiple bacterial and helminth infections^43^. Rare and unconventional T cell populations were also represented^19,61–63^. These included both 3’ and 5’ 10X chemistries, and a dataset generated by split-pool technology also performed well.

As an illustrative example, we applied T-RBI to a dataset from Miller et al. profiling gp33 tetramer–positive CD8⁺ T cells during chronic LCMV infection (day 28 post-infection)^64^. The original study identified five transcriptionally distinct clusters (**Fig. 5a**). Following T-RBI mapping, these cells projected coherently onto the CD8 MDE while preserving separation of the original clusters (**Fig. 5b**). Importantly, immgenT-based annotation further subdivided these cells into multiple immgenT clusters, revealing a richer organization of CD8⁺ T cell states than initially described (**Fig. 5c**). Only 5% cells from this dataset could not be confidently annotated (**Fig. 5d**). These unassigned cells were dispersed across the original UMAP, rather than belonging to a defined cluster (**Extended Data Fig. 6a**). This indicated that T-RBI’s hesitation at confidently attributing cells to an immgenT cluster likely reflected transitional or boundary states rather than novel T cell states absent from the immgenT dataset.

**Figure 5.**
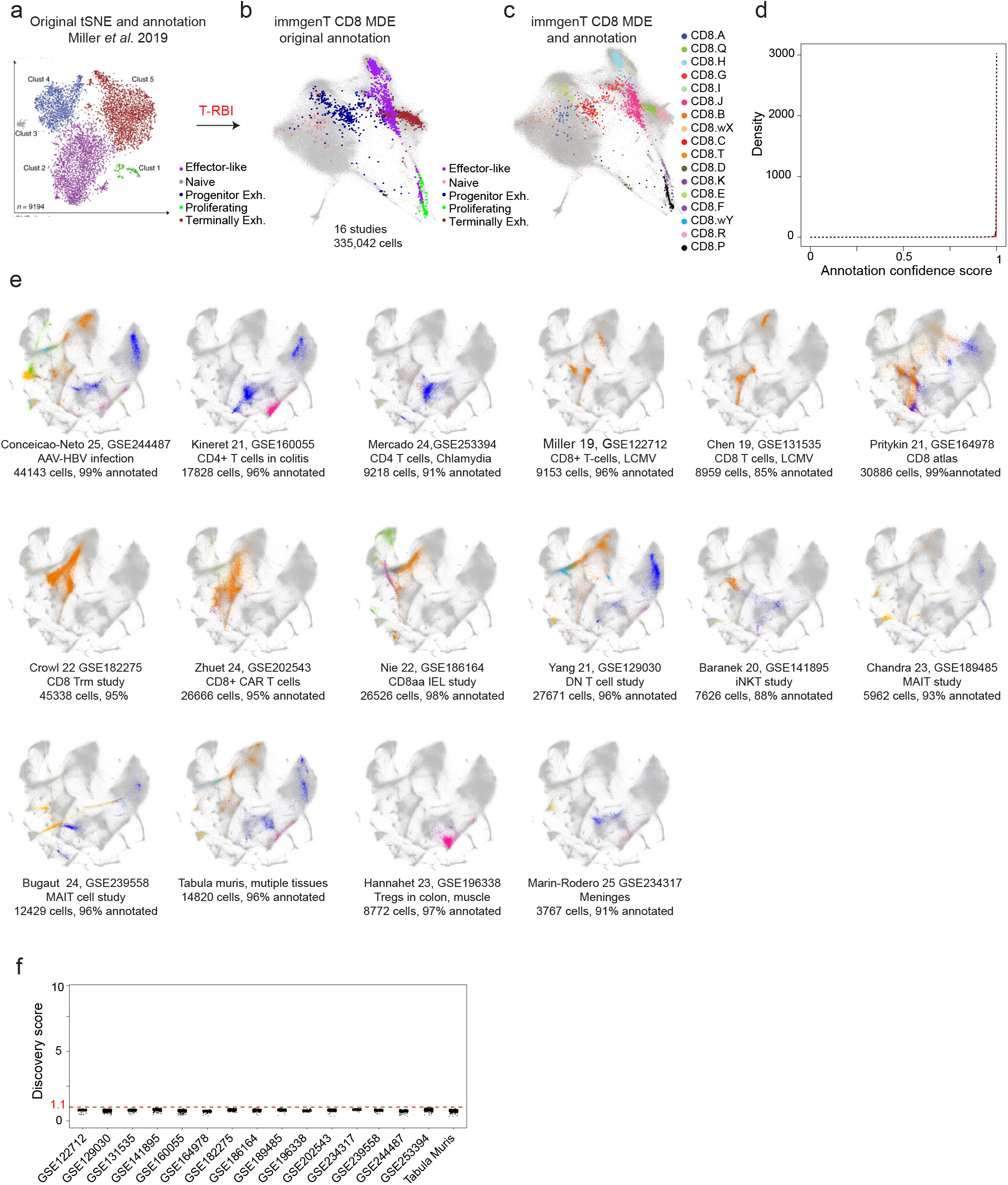
T-RBI integrates external datasets into the immgenT framework and supports near-saturation of T-cell states. (**a–d**) Example integration of an external study by Miller *et al.*^64^, which profiled GP33 tetramer–positive CD8⁺ T cells during chronic LCMV infection (day 28 post-infection). **(a)** Author-reported tSNE embedding with original cluster assignments and annotations. **(b)** Same query cells after integration with T-RBI, projected into the immgenT CD8-specific MDE and colored by the author’s original clusters. **(c)** Query cells in the CD8-MDE colored by immgenT cluster annotations. **(d)** Distribution of scANVI confidence scores for immgenT cluster assignments of query cells. **(e)** Gallery of 16 external datasets (**Extended Data Table 5**) mapped to the immgenT reference. For each dataset, query cells are highlighted within the all-T MDE and colored by lineage. Numbers indicate total cells mapped and the percentage annotated at the cluster level. **(f)** Distribution of discovery scores (see **Methods**) across external studies. The discovery score estimates whether query T cells form coherent states outside the immgenT reference. A threshold of > 1.1 (conservative) indicates T cells with potential novel states.

The broader results are depicted in **Fig. 5e**. As expected, contaminating non–T cells present in these datasets failed to map to immgenT and localized to distinct regions of the embedding (**Extended Data Fig. 6b**). In contrast, more than 99% of T cells were confidently assigned to level1 lineages, with concordant expression of canonical markers including *Cd4*, *Cd8a*, *Cd8b1*, *Foxp3*, *Zbtb16*, and *Trdc* (**Extended Data Fig. 6c,d**). At finer resolution, most query T cells (mean 97%, range 85-99%) received confident level 2 cluster assignments (**Fig. 5e, Extended Data Fig. 6e**). The remaining unannotated cells embedded within regions populated by annotated T cells but were not assigned to discrete clusters, consistent with cells occupying intermediate transcriptional states rather than novel cell clusters (**Extended Data Fig. 6f**).

To formally distinguish transitional cells from genuinely novel T cell states, we devised a discovery score that quantifies whether unannotated cells preferentially associate with other unannotated cells. This score, defined as the ratio of k-nearest-neighbor distances between unannotated and annotated cells, was computed in the query dataset’s native PCA space, independently of immgenT integration, providing a conservative and reference-agnostic test for novelty. Across all 16 studies, unannotated T cells exhibited low discovery scores (<1.1, i.e., no more than 10% closer to each other than to annotated cells) (**Fig. 5f, Extended Data Fig. 6g**). As positive controls, non-T cells clustered together and away from annotated T cells, yielding high discovery scores (**Extended Data Fig. 6b,g,h**).

To assess whether the T-RBI procedure could identify novel T cell states as distinct, rather than forcing them into existing embeddings and clusters, we performed ablation tests (see **Methods**), in which individual reference clusters were iteratively removed. Cells from the omitted clusters mapped to the corresponding “holes” in the reference embedding (as expected from their transcriptional proximity with surrounding clusters and the local-distance optimization of MDE) (**Extended Data Fig. 7a)**. Many of these cells remained unannotated, whereas others were assigned to close neighboring clusters with reduced confidence scores (**Extended Data Fig. 7b,c)**. Their discovery scores were also increased, with a substantial fraction exceeding the 1.1 threshold of putative novel states (**Extended Data Fig. 7d)**.

Together, these analyses demonstrate that T-RBI enables robust integration of external datasets into the immgenT framework while preserving sensitivity to novel states (**Extended Data Note 4**). At the same time, the presence of unoccupied regions within the immgenT state space underscores the value of immgenT as a systematically designed reference dataset that extends beyond what can be achieved by aggregating existing datasets alone. T-RBI addresses a long-standing limitation in single-cell analysis whereby each new dataset is visualized in its own, non-transportable, dimensionality reduction map. By projecting query cells to a fixed MDE reference, T-RBI standardizes embeddings and enables direct, quantitative comparisons across studies. Importantly, T-RBI does not force alignment of query cells to existing states: query cells may occupy novel regions of the embedding, preserving sensitivity to previously unobserved T cell states.

### Distribution of shared T-cell states across tissues and perturbations

Using the immgenT framework, we examined how clusters are distributed across tissues and are remodeled during immunological perturbations. First, integration preserved expected differences in T cell composition across organs (**Fig. 6a,b, Extended Data Fig. 8**), demonstrating that datasets were aligned while retaining biologically meaningful variation. With the exception of gdT.R, corresponding to dendritic epidermal T cells (DETCs)^18^ in the skin, most clusters were detected across multiple tissues but at a broad range of frequencies. Blood, lymph nodes, and spleen showed similar compositions, enriched in naive-like CD62L⁺ clusters. However, resting states were not restricted to lymphoid organs, as CD8.B and CD4.B were preferentially represented in non-lymphoid tissues. In contrast, the small intestine and colon exhibited pronounced enrichment of CD8αα and γδ T-cell populations. Non-lymphoid organs, including kidney, liver, lung, and skin, were enriched for activated CD44⁺ T-cell states (most prominently CD8.Q, CD4.Q, and Treg.D/E) with this shift particularly marked in the skin, whereas other organs, such as the mammary gland, displayed more distinct compositional profiles. Non-conventional lineages also exhibited structured yet overlapping tissue distributions, like gdT.G/K. vs gdT.R/S/D/I, CD8aa.B vs CD8aa.A in the gut.

**Figure 6.**
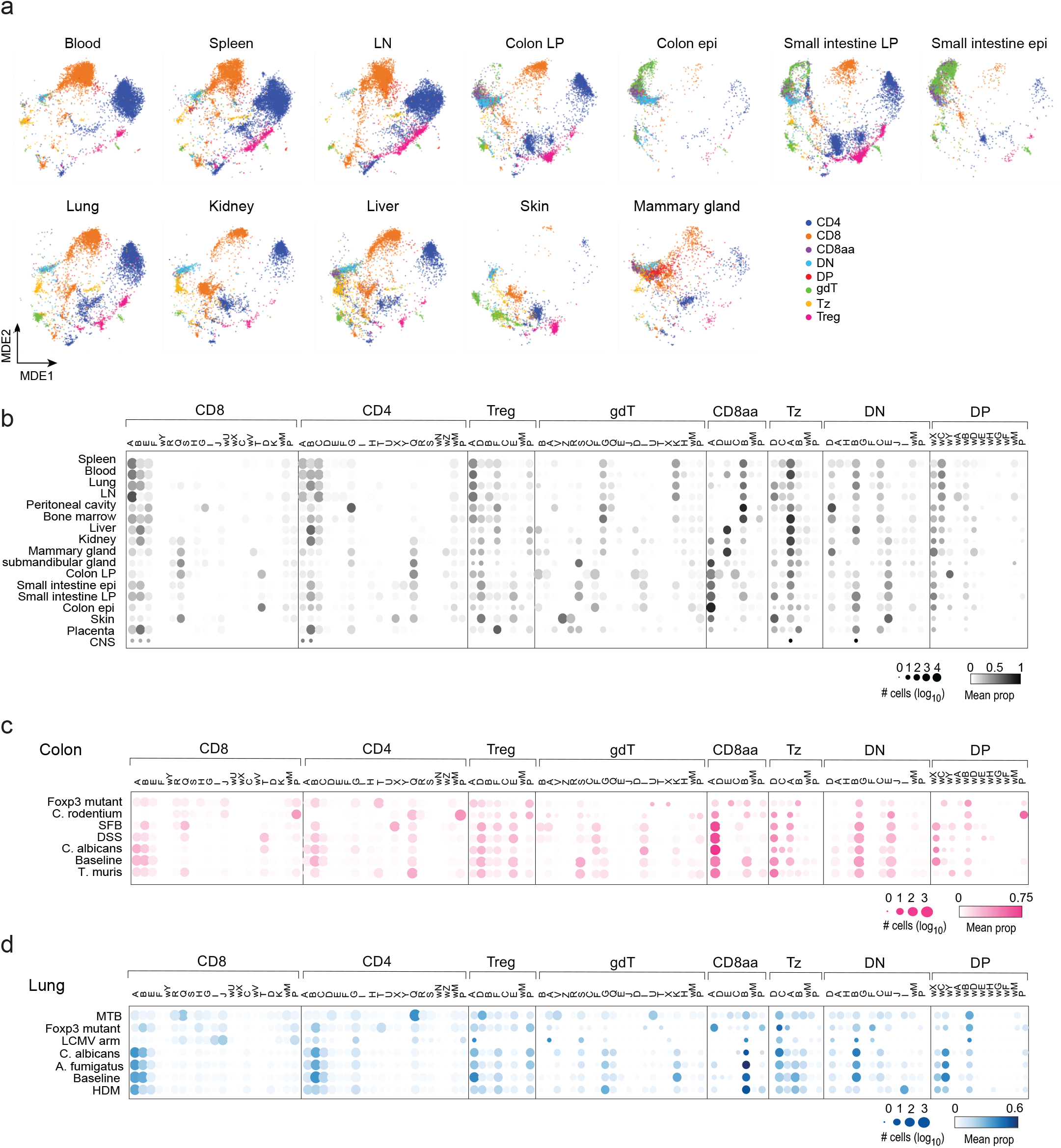
Reuse and redistribution of shared T-cell states across tissues and immune perturbations. **(a)** Gallery of all-T MDE showing the distribution of T cells at baseline across different organs, colored by lineage. **(b-d)** Dot plot showing the proportion of immgenT clusters across organs at baseline **(b)**, across immune perturbations in the colon **(c)** and across immune perturbations in the lung **(d).** Color intensity reflects the mean proportion of each cluster within each lineage, and dot size indicates the total number of profiled cells.

Cluster distributions were further remodeled during immune challenges in the colon and lung, revealing shared principles of state redistribution across tissues (**Fig. 6c,d**). In both organs, CD8.Q cluster was enriched during infection and inflammation. CD4.Q (IFNγ-producing, multipotential) was broadly induced across perturbations, whereas other CD4⁺ clusters showed more context-specific responses, including enrichment of the IL-5–producing CD4.T in *Foxp3*-deficient mice, and the IL-17–producing CD4.X specifically during SFB colonization. *Citrobacter rodentium* infection induced widespread proliferation across CD4, CD8, regulatory T cells, and γδ T cells, whereas *Candida albicans* colonization resulted in minimal deviation from baseline in both tissues.

Together, these results show that the immgenT framework captures tissue-and perturbation-associated variation within a unified organizational framework for T-cell states. Rather than generating new cell identities, diverse immunological parameters (including tissue, perturbation, time, antigen specificity, and ontogeny) primarily act by redistributing and differentially enriching a shared set of T-cell states.

### Effector gene expression reflects both cluster identity and immune context

Effector molecules, including cytokines, chemokines, secreted enzymes, and surface-bound ligands, enable the palette of T cell functions^65–68^. Textbook examples include cytokines that guide immunoglobulin class switching in B cells, IL-10 and TGFβ that mediate inhibitory functions, or mediators of target cell death. These associations have often been treated as defining features of T-cell identity, leading to simplified models in which effector programs were considered binary, mutually exclusive, and organized around antagonistic cytokine pairs^69^. However, the breadth of the immgenT data brought a comprehensive perspective on effector molecule deployment across lineages, states, and locations that had been lacking. We systematically examined the expression of T cell effector genes selected by Gene Ontology and expert curation (**Fig. 7a**; other relevant molecules in **Extended Data Fig. 9a**).

**Figure 7.**
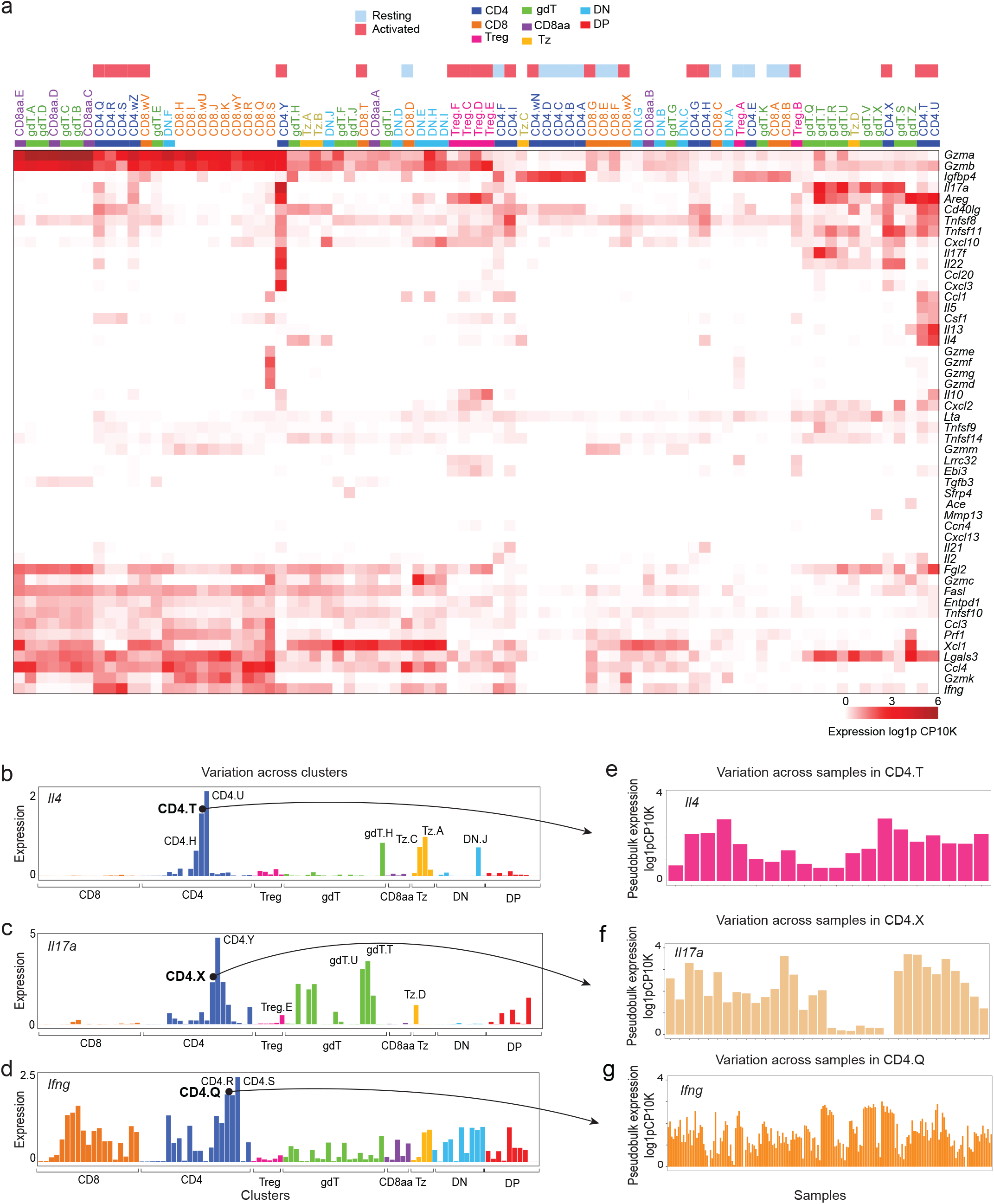
Effector gene expression across T cell states. **(a)** Heatmap of curated effector gene expression across immgenT clusters (log1p CP10K). Rows (genes) and columns (clusters) are ordered by hierarchical clustering. Annotation bars indicate lineage and activation status (resting versus activated). **(b–d)** Bar plots showing average expression of *Il4* **(b),** *Il17a* **(c)**, and *Ifng* **(d)** across immgenT clusters. **(e–g)** Bar plots illustrating sample-to-sample variability of effector gene expression within selected clusters: *Il4* expression in CD4.T **(e),** *Il17a* expression in CD4.X **(f)**, and *Ifng* expression in CD4.Q **(g)**.

In general, the expression of these effector functions was tied to cell activation, as the CD62L^+^ resting clusters (e.g. CD4.A/B and CD8.A/B) minimally express these molecules **Fig. 7a**. These cells were not completely inert, however, and did express *Tnf, Lta*, *Cd40lg*, at levels often not dissimilar from activated clusters, and even some *Gzma*. Within activated clusters, patterns of key cytokine expression did not follow rigid delineation, as illustrated by Th2 cytokines. *Il4*, *Il5* and *Il13* were all enriched in CD4.T/U, consistent with classical Th2 associations. But while *Il5* was very specific to CD4.T/U, *Il13* was also detected in γδ T-cell cluster gdT.Z, and *Il4* exhibited an even broader spread of expression (**Fig. 7a,b**), extending to other γδ T-cell clusters (gdT.H), Tz-cell clusters (Tz.C and Tz.A), and CD4.H, a cluster enriched for Tfh-like cells that also expressed *Il21*. Thus, beyond the main Th2-like node of CD4.T/U, several other clusters variably expressed Th2 cytokines, in keeping with ref^70^. Within the IL-17 family, *Il17f* showed greater specificity than *Il17a*, only partially overlapping across CD4, gdT, and Tz clusters (**Fig. 7a,c**). *Il22* largely overlapped with *Il17a/f*-expressing clusters, consistent with partial co-regulation. *Il9* was lowly detected in CD4 and Tregs (**Extended Data Fig. 9b**). But the greatest departure from the canonical one-cytokine/one-cell-state model was perhaps for *Ifng*, whose expression was particularly widespread (**Fig. 7a,d**). As expected, *Ifng* was absent from resting CD4 and CD8 clusters, but it was expressed in a great number of CD4, CD8, γδ T, CD8αα, and Tz clusters.

With nine members, the granzyme family provided another illustration of effector diversity. Traditionally considered terminal executioners in different facets of cytotoxicity, granzymes also influence other immunological processes in a less dramatic fashion, including antigen presentation and cytokine activation^71^. Granzyme expression partitioned in interesting patterns (**Fig. 7a**). *Gzma* and *Gzmb* were strongly and broadly expressed in many activated clusters of CD8, CD8αα, and γδ T cells (associated with other death effectors like *Prf1* and *Fasl*, and more loosely with *Ifng*), but also at lower levels in CD4 and Treg cells. *Gzmk* and *Gzmc* followed the same trend, but with more specific profiles. A striking group (*Gzmd/e/f/g*) was sharply restricted to CD8.S and CD4.Q, and *Gzmm* was another outlier. Thus, granzymes organize the space of T cell diversity along yet other axes.

Importantly, mean expression across clusters captured only part of the underlying variability. Even in prototypical clusters, such as CD4.Q for *Ifng*, CD4.T/U for *Il4*, or CD4.X/Y for *Il17a*, expression varied across a wide range across samples (5-to 40-fold, **Fig. 7e-g**), indicating that the expression of effector molecules is affected by other contextual influences, and not merely by cluster identity alone.

Together, these results reveal a richer, more nuanced landscape of effector-molecule expression than classical models suggest. Even among effector genes selected for high cluster specificity, expression patterns were broader than any well-defined subset(s), and further shaped by context. These findings caution against simple deductive interpretations—for example, inferring cell identity solely from the presence of a single cytokine—and instead emphasize that T-cell states are defined by complex combinations of effector programs whose deployment depends on both molecular state and immune context.

### Transcription factor expression across the T cell clusters

Transcription factors (TFs) are key regulators of T cell differentiation, function, and stability. In rare cases, TF expression is specific to a defined T cell lineage or state (e.g. FOXP3 in Treg cells), but the relationship between TF expression and T cell identity and function is generally more complex^72^. It was thus of interest to systematically analyze TF expression across immgenT lineages and clusters. First, we generated a receiver operating characteristic (ROC)-like graph with an area under the curve (AUC) score that sums each TF’s cumulative expression across ranked clusters (**see Methods, Fig. 8a,b**). At the lineage level, very few TFs were lineage-specific, with standouts including *Foxp3* in Tregs and *Zbtb16* in unconventional Tz cells (**Fig. 8a, Extended Data Fig. 10a, Extended Data Table 6**). At cluster resolution, several other TFs emerged (**Fig. 8b, Extended Data Fig. 10b,** individual profiles in **Fig. 8c-m, Extended Data Fig. 10c-f, Extended Data Table 7**), *Sox13* had highly specific expression in a subset of gdT cell clusters (**Fig. 8e**), as expected^73^. *Rorc* was also restricted to a few defined clusters across multiple lineages, including CD4 T cells (CD4.X/Y), Tregs (Treg.E), and subsets of gdT cells and Tz cells (**Fig. 8f**), strikingly concordant with *Sox13* in gdT cells, consistent with Malhotra et al.^74^. This analysis also revealed TFs with narrow expression profiles in Th2-like CD4.T/U, such *Pparg*^75^ or *Hes1,* strongly enriched with greater specificity than *Gata3* in this context (**Fig. 8g,h,i**). Other intriguing TFs that have received limited attention in T cells, like *Sox4*, *Ztbt32*^76^, *Tshz2*^77^, or *Hlf*^78^, were also highlighted by this specificity analysis (**Extended Data Fig. 10c-f**).

**Figure 8.**
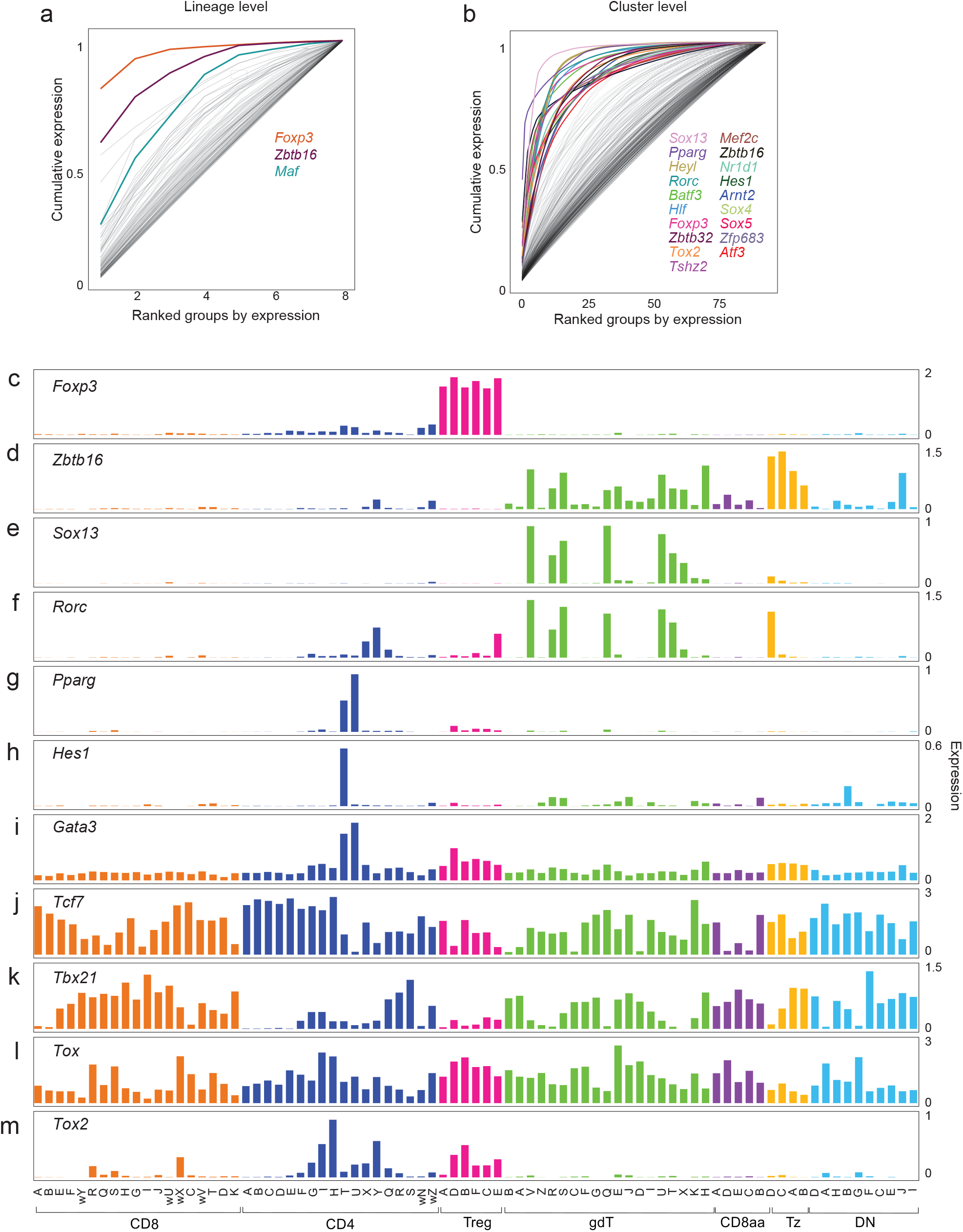
Transcription factor expression across T-cell states. **(a)** ROC-like plot of transcription factor (TF) specificity across T-cell lineages (see also **Extended Data Table 6**), showing normalized cumulative TF expression across ranked lineages. The three most lineage-specific TFs are highlighted. **(b)** Same analysis as in (a), performed across immgenT clusters. The topmost cluster-specific TFs are highlighted and listed in decreasing order of specificity (see also **Extended Data Table 7**). **(c-m)** Bar plots showing expression of selected TFs across immgenT clusters: *Foxp3* **(c)**, *Zbtb16* **(d),** *Sox13* **(e),** *Rorc* **(f),** *Pparg* **(g)**, *Hes1* **(h)**, *Gata3* **(i)**, *Tcf7* **(j)**, *Tbx21* **(k)**, *Tox* **(l)**, and *Tox2* **(m).**

In contrast, patterns for several TFs were less congruent with classical expectations. *Gata3* was broadly expressed, with only relative over-expression in CD4.T/U (**Fig. 8i**). *Tcf7* (**Fig. 8j**) or *Tbx21* (**Fig. 8k**, which encodes Tbet) were not confined to discrete populations within any lineage. Tbet is often thought to drive specific programs within discrete subsets of several lineages (e.g., Th1, NKT1^79^), but it was present in most activated clusters. Among TFs associated with exhaustion, *Tox* is often represented as a “typical exhaustion factor”^80^, but was broadly expressed across many clusters, with only modest enrichment in CD8.R/S as might have been expected (**Fig. 8l**); *Tox2* showed more specificity, but highest in CD4 T cells and Tregs, not CD8 T cells (**Fig. 8m**).

Together, these results reveal a rich and nuanced TF expression landscape in T cells that supports some canonical associations but not others, and highlight previously underappreciated TF–state relationships.

### T-cell states as combinations of gene programs

Analyses of TFs and effector molecules caution against inferring T-cell identity from the presence or absence of any single transcript. Instead, their shared, quantitative expression across clusters suggests that T-cell states are best defined by combinations of gene programs (GPs), in line with previously proposed models^81–84^. To formalize this idea, we performed GP analysis across the immgenT data, detailed in a companion manuscript (**immgenT-GP**), combining two complementary approaches. Briefly, we first used empirical Bayes matrix factorization^85^ to embed both genes and cells into a common, interpretable GP space, where each cell’s transcriptome is represented as a weighted combination of gene programs, placing cells along continuous axes of program activity rather than in discrete categories (**Fig. 9a**). Complementarily, we developed a variational autoencoder-based framework that iteratively learns a discrete regulatory codebook that orthogonally validated the matrix factorization structures.

**Figure 9.**
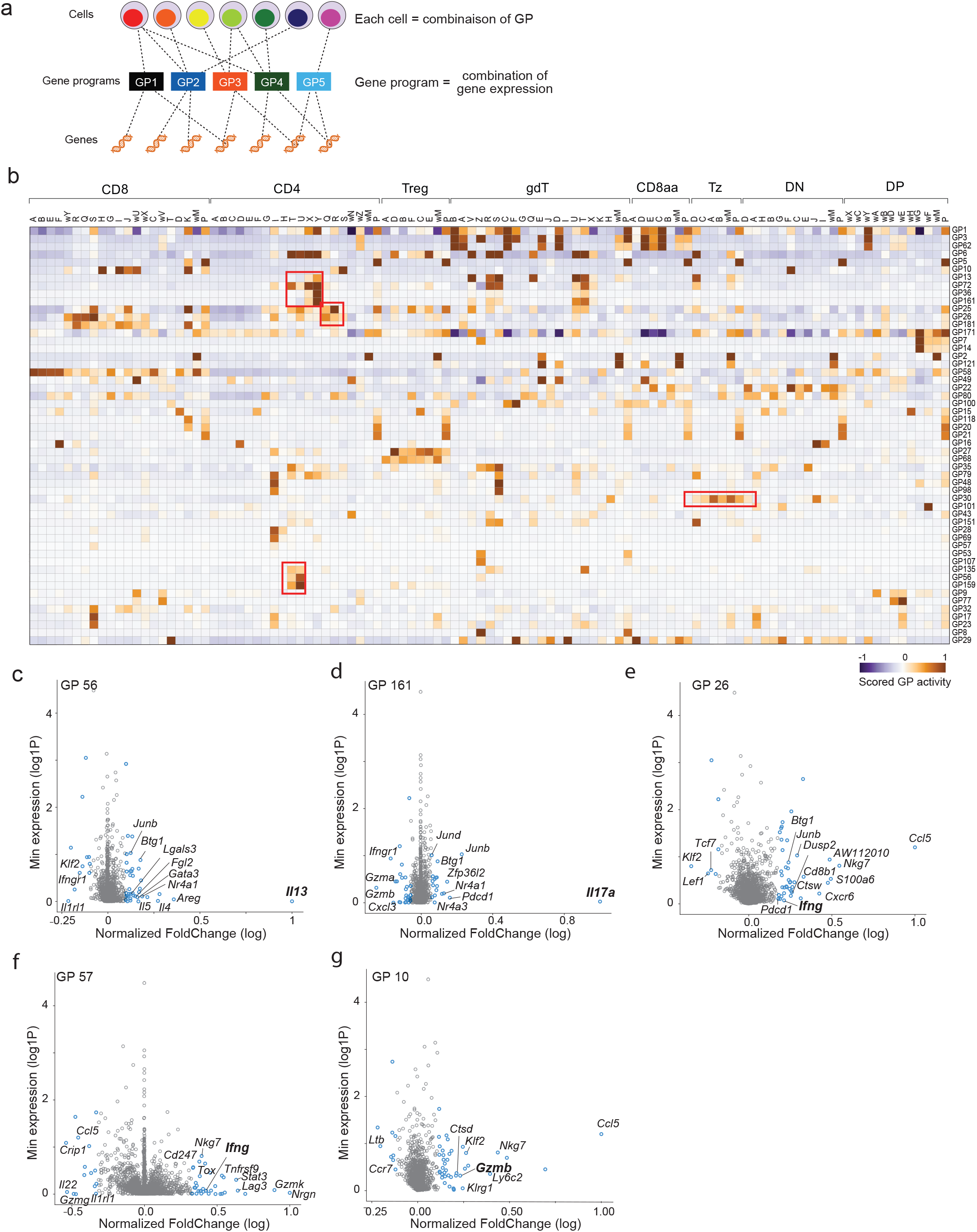
T-cell states are defined by combinations of gene programs. **(a)** Schematic of the gene-program (GP) analysis illustrating shared gene programs across cells and clusters. Genes are grouped into a limited number of transcriptional programs (GP), and each cell’s gene-expression profile is represented as a weighted combination of these programs. **(b)** Dominant gene programs using empirical Bayes matrix factorization across all immgenT. The heatmap shows scaled GP activity, represented as effect sizes from one-versus-all linear models for each cluster. Gene programs were selected based on an effect size > 0.5 in at least one cluster. **(c–g)** Mean expression versus fold-change plots for representative gene programs: GP56 (Th2-like) **(c),** GP161 (Th17-like) **(d)**, GP26 (Th1-like) **(e)**, GP57 (Th1-like) **(f),** and GP10 (Cytotoxic-like) **(g).** Values are log-normalized, and fold changes are scaled relative to the maximal induction observed for each gene program.

GP inference was performed independently of totalVI integration of the cells into clusters, but the expression of the programs reflected well the overall structure of the atlas (**Fig. 9b**), independently validating its relevance. This process condensed expression patterns of over 20,000 genes into fewer than 200 robust GPs, providing a compact representation of transcriptional heterogeneity. With a few exceptions, clusters were not defined by unique programs; each state was characterized by a distinctive combination of shared programs.

Several intuitive examples illustrate this principle. GP30 and GP68 were strongly enriched in Tz and Treg cells (**Fig. 9b)**, respectively, mirroring the lineage-specific transcription factors highlighted in earlier analyses. Effector functions were deconstructed into multiple overlapping programs. GP56, with *Il13* as a standout, was strongly expressed in CD4.T and CD4.U, but also in gdT clusters (**Fig. 9b,c)**. Yet GP56 co-occurred with additional programs (GP135, GP159) (**Fig. 9b**), each showing distinct distributions across clusters, together simplifying the complex expression patterns of *Il4*, *Il5*, and *Il13* described above. *Il17a* expression was captured primarily by GP161 (**Fig. 9b,d**). IFN-γ expression, which appeared promiscuous in **Fig. 7a**, was partitioned into multiple programs, mainly GP26 and GP57 (**Fig. 9b,e,f**). These programs shared interferon-responsive genes (e.g., *Stat3*, *Ifngr1*) but differed in their co-expressed genes (e.g., *Ccl5* and *Gzmk*) and biological context: GP26 represented a broadly activated interferon program across CD8 T cells, whereas GP57 selectively tuned IFN-γ–associated expression in CD8.S, a cluster enriched in cancer. Cytotoxicity-associated transcripts such as *Gzma* and *Gzmb* were driven by GP10 (**Fig. 9b,g**), whose activity extended to clusters of gdT, CD4 and CD8aa cells, illustrating how shared programs span lineages while contributing to cluster-specific phenotypes.

The gene-program framework complements immgenT cell cosmology by reducing complex transcriptomes to a limited set of interpretable programs. This representation enables a quantitative, continuous description of T-cell states, clarifying the combinatorial logic underlying effector functions.

### Beyond markers and signatures: integrative definition of T cell states

A long-standing quest in immunology has been to define T cell subsets and associate them with specific functions (e.g., helper, cytotoxic) in one-to-one relationships. Successive technical shifts have shaped this goal, each expanding the granularity of possible definitions. Early frameworks relied on surface markers and increasingly complex combinations thereof, followed by transcriptomic approaches that introduced gene signatures as higher-dimensional alternatives. Here, we re-examined these concepts within the immgenT framework, comparing the performance of simple marker combinations, gene signatures, and integrative mapping approaches.

As a concrete example, we discuss tissue-resident memory CD8^+^ T cells (T_RM_). Although residency and memory are cellular properties not directly tested here, we operationally define T_RM_ as LCMV–specific (P14 transgenic) CD8^+^ T cells located in non-lymphoid tissues at memory timepoints (30 days post-infection (dpi), after antigen clearance, hereafter P14d30), a population extensively profiled in IGT36/38/40, including in the small intestine^86^. These are unambiguously mapped to CD8.Q (**Fig. 10a**). The combination of CD103 (*Itgae*) and CD69 in flow cytometry is classically used to evaluate or purify T_RM_ cells, although it has been questioned as closer examination reveals substantial heterogeneity^54,87^. In the immgenT data, the CD103^++^CD69^+^ criterion identifies many other cells, in other activated clusters and even in the naive group (**Fig. 10b**). Reciprocally, the IGT38 CITEseq data show that the CD103/CD69 profile of P14d30 T_RM_ cells varies with the parenchymal tissue in which they are located (**Fig. 10c**): CD103 is high in the intestine, uneven in the prostate, and missing from half of P14d30 cells in the salivary gland, in keeping with Crowl et al.^54^. Thus, CD103 is neither fully specific nor fully sensitive for T_RM_, and CD8.Q includes T_RM_ that are both CD103^+^ and CD103^-^, illustrating the complex combination of protein expression defining cell states.

**Figure 10.**
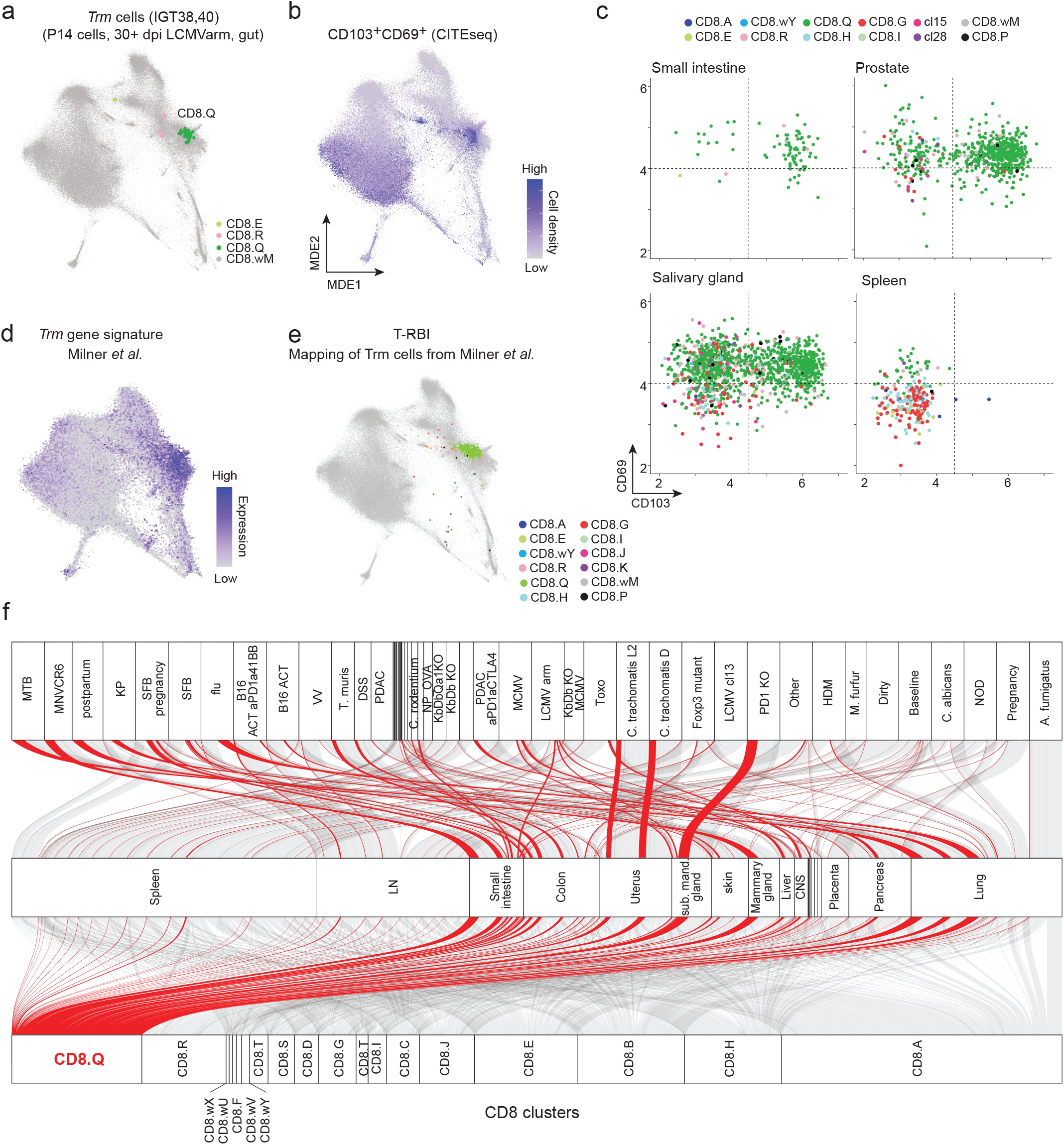
AI-based reference mapping reveals the limits of marker-and signature-defined T-cell identity. **(a)** CD8-MDE highlighting resident-memory CD8⁺ T cells (Trm) mapping predominantly to CD8.Q. Trm cells are defined here as LCMV-specific (P14) CD8⁺ T cells isolated 30 days after LCMV Armstrong infection in the gut. Cells are colored by immgenT cluster. **(b)** CD8-MDE with CD103⁺CD69⁺ T cells in the full immgenT dataset extending beyond CD8.Q, indicating that canonical Trm markers do not uniquely define this molecular state. The gating strategy is shown in (c). **(c)** CD69 versus CD103 expression in memory cells from the small intestine, prostate, salivary gland, and spleen (CITEseq, log1p(CP10K); P14 CD8⁺, 30 dpi), illustrating that canonical Trm markers are variably expressed across tissues, including within CD8.Q. Cells are colored by immgenT clusters. **(d)** CD8-MDE with expression of a published Trm gene signature across all immgenT cells (signature from Milner *et al.*^89^), showing that Trm signature, like markers, does not uniquely define CD8.Q. **(e)** T-RBI mapping of Trm cells from Milner *et al.*^89^ (the same cells used to derive the Trm signature shown in (d)) map unambiguously into CD8.Q. **(f)** Alluvial plot showing the contribution of CD8.Q cells (red) across a broad range of organs and immune perturbations, illustrating CD8.Q as a transcriptional basin into which cells from diverse biological conditions converge.

**Figure 11.**
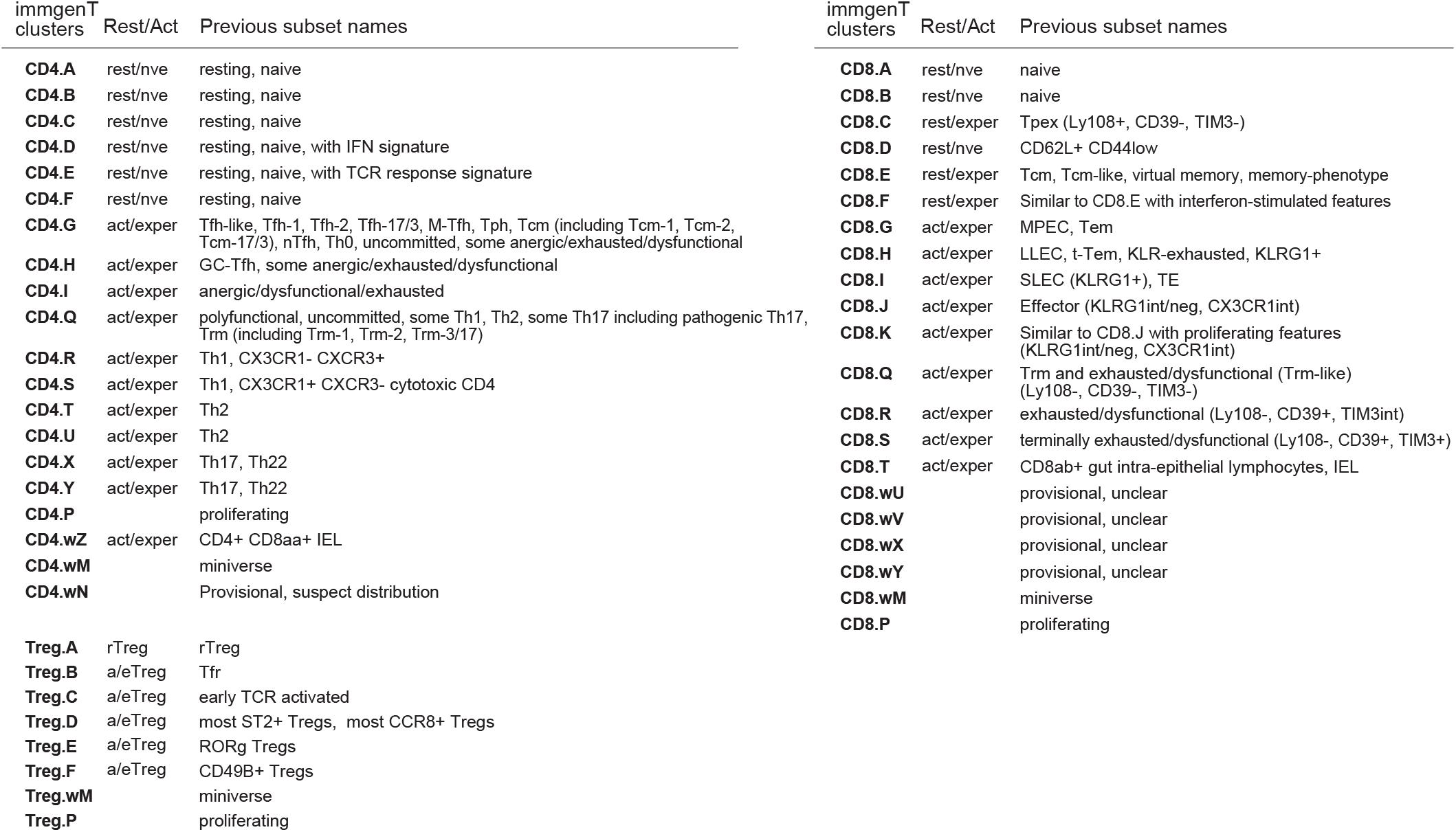

Another illustration of the limitation of single-marker–based definitions of T cell states comes from the expression of commonly used identifiers of Th differentiation (**Extended Data Fig. 11a,b**). Neither *Cxcr3* or *Il12rb2* (Th1), *Ccr4* or *Il4ra* (Th2), and *Ccr6* (Th17) were in any way specific for the polarized clusters of CD4^+^ T cells (*Il23r* in Th17 tip the sole exception). There is a caveat here, in that we are measuring RNA levels, and translational control may make protein expression more restricted. Similarly, relying on TFs like *Tox* or *Tcf7* (encodes TCF-1) to identify exhausted or “stemness” states is highly problematic, given their widespread expression (**Fig. 8j,l,m**). While *Tox* or *Tcf7* are demonstrably necessary for exhausted states or T cell renewal properties^80,88^, their presence is not sufficient to infer that all *Tcf7*-or *Tox*-positive cells have stem or exhausted characteristics, a frequent misrepresentation.

The advent of whole-genome transcriptomics suggested that expanding from single markers to multi-gene signatures might overcome the limitation of low-marker definitions. We therefore evaluated published T_RM_ gene signatures, including those derived by Milner et al.^89^ As shown in **Fig. 10d**, this signature is enriched in regions that include CD8.Q and P14d30 cells, yet it clearly lacks specificity (as a caveat, the signature index is continuous, whereas the cell attributes are categorical, potentially making the comparison unfair). Similarly, a sharply defined core Treg signature^90^ is strongly enriched in Tregs relative to most CD4+ cells, but is also high elsewhere (**Extended Data Fig. 11c**). Thus, gene signatures have limited value for unambiguous cell-state definition. In considering why signatures perform poorly, several points became apparent. First, signatures are gene lists defined by arbitrary thresholds (e.g., fold change, p-value) and typically lose quantitative information (all genes weighted equally, irrespective of degree of specificity). Second, and most importantly, signatures are only valid in the specific context in which they were defined, comparing one target to a reference population (typically A vs. B, or A vs. all other subsets). One can clearly define the target (e.g., T_RM_, Treg), but the reference population changes. There is no intrinsic, and cell-autonomous “Treg signature”. While signatures have value as part of large collections in providing color to a gene set^91^, they are insufficient (and possibly misleading) to annotate cell clusters within a single-cell dataset.

Here, and building on the results of Figure 5, we propose a new strategy for cell identification that does not rely on markers or signatures but on AI embedding into a reference dataset. The cells originally used to define the T_RM_ signature of **Fig. 10d** mapped unambiguously to CD8.Q with high sensitivity and specificity (**Fig. 10e**), demonstrating that integrative, reference-based approaches outperform both single markers and gene signatures for cell-state definition. This improvement arises because T-RBI captures high-dimensional and nonlinear relationships across the transcriptome, including conditional dependencies between genes and heterogeneous expression patterns across subsets of cells, rather than relying on linear combinations of a restricted gene list. Importantly, CD8.Q was not specific to LCMV, but was a convergent T cell state observed in many other infections in NLT at memory timepoints (**Fig. 10f**), including Chlamydia-specific T cells in the uterus or flu-specific T cells in the lung. In those contexts and tissues where markers and gene signatures defined in LCMV might not translate well, T-RBI integration into the immgenT framework remained efficient.

We therefore propose T-RBI as a more accurate and generalizable framework to define T cell states. As discussed in **Extended Data Note 2**, ImmGen now provides a web interface for public use.

## Discussion

immgenT is a community effort that started with two ambitious goals: to comprehensively profile mouse T cells across organs and immunologic perturbations, and to determine whether T cell states can be organized within one coherent molecular-based framework^27^. Despite, but also because of the breadth of the data, a comprehensive picture does emerge: T cell diversity occupies a finite and structured landscape rather than an ever-expanding universe of discrete identities. The 107 AI-resolved clusters form robust and reproducible landmarks that recur across tissues and challenges, establishing the basis for a unified molecular classification of T cells across the mouse body.

Why a molecular reference for T cells? Historically, T-cell classification has relied on a heterogeneous mix of criteria, including cytokine production, surface-marker expression, developmental origin, tissue localization, inferred function (helper, cytotoxic), and cell experience (naive, memory). While each of these dimensions captures meaningful biological properties, together they form an eclectic taxonomy, shaped as much by available assays as by underlying biology. Such classifications can obscure unrecognized or shared functions, overlook quantitative variation, and implicitly assume that a particular function or a marker fully define cellular identity^92,93^. In contrast, the immgenT project arrives at a molecular reference that is robust because grounded in RNA and protein expression, and that appears comprehensive when confronted with external datasets. This notion of replacing a strict cell taxonomy based on a few markers with an AI oracle that returns assignments based on whole-genome activity is also where the Human Cell Atlas gravitated to^94^. Along this line, we are actively investigating how faithfully the immgenT framework translates to human T cells.

Hopefully, immgenT will not merely be a static reference, published and then forgotten; through public T-RBI (**Extended Data Note 4**) and the Rosetta web applications (**Extended Data Note 2**), this molecular reference is directly accessible, reusable, and directly connected to experimental immunology. These web resources also support re-usability in two complementary ways: the CITE-seq data in Rosetta enables cytometry-based measurements to be positioned within the full molecular landscape; the computation run by T-RBI allows single-cell transcriptomic datasets to be mapped unambiguously onto the immgenT reference space. In addition, ImmGen will evolve the framework over time: T-RBI users are asked if the consortium can retain their submitted data, for incorporation into a growing “crowd-sourced” reference. We should acknowledge limitations to the framework proposed here. For one, the cluster solution provides a discrete approximation of the continuum of T cell states and will likely evolve in the future, particularly around the provisional clusters, much as releases of genome sequences do. Secondly, future data that are deeper and/or selectively focused on a region of the T cell space, as well as computational refinements, may reveal more fine-grained structures within the currently defined states.

immgenT’s unified molecular landscape helps clarify common conceptual pitfalls in the study of T-cell identity. One frequent syllogism equates the expression of one functional transcript with identity. For example, simply because *Tox* expression is necessary for “exhausted” functionality, inferring that all *Tox-*positive cells are exhausted. Or that because IFNγ production occurs in a “Th1” state, any IFNγ-expressing CD4+ T cell must be a Th1. While IFNγ-expressing T cells are enriched in several CD4 clusters (Q/R/S), its expression is broader, and additional molecular context is required to precisely define the cells under study. Function-based classifications introduce a second common misconception: equating function with molecular identity. Concepts such as tissue residency, memory, exhaustion, effector, or helper specialization are often treated as intrinsic cell identities. In the immgenT framework, however, such functions map onto multiple molecular states and show substantial overlap. Conversely, individual immgenT states often contain cells drawn from diverse immunological contexts, only some of which display a given function. This decoupling highlights the relationship between state and function. We propose that many T-cell functions are emergent properties of molecular states that are necessary but not sufficient on their own, requiring appropriate environmental context. For example, activated CXCR5⁺PD-1⁺ cells in cluster CD4.H encompass Tfh cells which can be found inside GCs, with specialized functions in helping B cell affinity maturation, but also outside of GCs with other functions^95,96^. Describing such cells by their molecular state (CD4.H) allows for explicit discussion of the cell *state*, independently of the location-dependent *function*. **Extended Data Table 8** and **Fig. 11** attempt to summarize the correspondence between immgenT clusters and previously named subsets or other common descriptors (acknowledging that this table will evolve with community input).

Each immgenT cluster can be viewed as a transcriptional attractor—a stable region of gene-expression space toward which cells converge across tissues and perturbations^97–99^. This convergence reflects the reuse of core gene programs, including transcription factors and effector modules, across many contexts. The presence of cells from multiple conditions within each state underscores the and pleiotropy of T-cell responses and suggests potential avenues for cell engineering or synthetic manipulation, once we establish the causal combination of TFs that drive a particular GP or cluster^72,100^. The GP framework further refines this view by reducing complex transcriptomes to a limited set of reusable building blocks. Future integration of genetic variation, epigenetic regulation, and post-transcriptional control will be necessary to fully explain how cells move between transcriptional basins and how stable or reversible these states are.

More than a change in terminology, immgenT provides a conceptual and practical foundation for studying T-cell heterogeneity. Focusing on molecular states should facilitate communication, enable more precise comparisons across studies, and lead to new hypotheses about how immune functions emerge from shared molecular programs.

## Supporting information

Extended Data Table 1 - Sample Metadata

Extended Data Table 2_citeseq_panel

Extended Data Table 3_QC_RNA_ADT_sample_summary_table

Extended Data Table 4_citeseq_qc

Extended Data Table 5_T-RBI

Extended Data Table 6_AUC_TF_lineage

Extended Data Table 7_AUC_TF_clusters

Extended Data Table 8 immgenT_PreviousNomenclature

## Acknowledgments

We thank the many colleagues who were consulted at various stages of this project. This work was funded by a grant from the NIH to the ImmGen consortium (R24-072073).

## Author Contributions

IM, OBR, DB, GB, SB, JC, EF, AF, GG, AMG, TH, RL, ZL, ALF, EL, JM, KO, SPR, TS, CT, AW, and SZ performed experiments, OC, SSP, LZ, MF, CEM, MF, CI, LY, MB and DZ participated in the computational processing and analyses, AZF and JC devised and contributed reagents; DZ, AG, MK, JK and CB designed the overall study; MLA, AB, RB, LB, KC, AC, LG, AH, JH, II, BJ, SJ, MK, SK, DK, VK, MN, MP, DS, SS, MS, MS, UvA and NY oversaw the experiments; IM, OBR, DB, GB, SB, EF, AF, GG, AMG, TH, CI, RL, ALF, EL, KO, SPR, TS, CT, EW, SZ, LG participated in analytical workshops; DZ and CB wrote the manuscript with input from other authors.

## Collaborators

Participants in the immgenT Project include: Aaron Liu¹, Alexander Chervonsky², Alexandra Cassano², Alia Welsh³, Amir Ferry⁴, Ananda Goldrath⁴, Andrea Lebron-Figueroa⁵, Ankit Malik², Anna-Maria Globig⁶, Antoine Freuchet², Bana Jabri², Charlotte Imianowski⁷, Christophe Benoist⁵, Claire Thefaine⁸, Dan Kaplan⁷, Dania Mallah⁵, Dario Vignali⁷, David Sinclair⁵, David Zemmour², Derek Bangs⁹, Domenic Abbondanza², Enxhi Ferraj¹⁰, Eric Weiss⁷, Erin Lucas⁸, Evelyn Chang¹⁰, Gavyn Chern Wei Bee¹¹, Giovanni Galletti⁴, Ian Magill⁵, Iliyan D. Iliev¹², Joonsoo Kang¹⁰, Jordan Voisine², Josh Choi⁵, Julia Merkenschlager¹³, Jun R. Huh⁵, Katharine Block⁸, Ken Cadwell¹¹, Kennidy K. Takehara⁴, Kevin Osum⁸, Laurent Brossay¹⁴, Laurent Gapin¹⁵, Liang Yang⁵, Lizzie Garcia-Rivera¹, Marc K. Jenkins⁸, Maria Brbic¹⁶, Maria-Luisa Alegre², Marion Pepper⁹, Mariya London¹⁷, Matthew Stephens², Maurizio Fiusco¹⁶, Melanie Vacchio³, Michael Starnbach⁵, Michel Nussenzweig¹³, Mitch Kronenberg¹⁸, Myriam Croze¹⁹, Nalat Siwapornchai⁵, Nathan Morris¹², Nicole E. Scharping⁴, Nika Abdollahi¹⁹, Nitya Mehrotra², Odhran Casey⁵, Olga Barreiro del Rio⁵, Paul Thomas²⁰, Peter Carbonetto², Remy Bosselut³, Rocky Lai¹⁰, Sam Behar¹⁰, Sam Borys¹⁴, Sara E. Hamilton⁸, Sara Mostafavi⁹, Sara Quon⁴, Serge Candéias²¹, Shanelle Reilly¹⁴, Shanshan Zhang⁵, Siba Smarak Panigrahi¹⁶, Sofia Kossida¹⁹, Stefan Muljo³, Stefan Schattgen²⁰, Stefani Spranger²², Steve Jameson⁸, Susan M. Kaech¹, Takato Kusakabe¹², Taylor Heim²², Tianze Wang⁹, Tomoyo Shinkawa¹⁰, Ulrich von Andrian⁵, Val Piekarsa⁵, Véronique Giudicelli¹⁹, Vijay Kuchroo⁵, Woan-Yu Lin¹², Ziang Zhang²

1. NOMIS Center, Salk Institute for Biological Sciences, 2. The University of Chicago, 3. National Institutes of Health, 4. University of California San Diego, 5. Harvard Medical School, 6. Allen Institute for Immunology, 7. Dept of Dermatology and Immunology, University of Pittsburgh, 8. University of Minnesota, 9. University of Washington, 10. UMass Chan Medical School, 11. University of Pennsylvania, 12. Weill Cornell Medicine, 13. The Rockefeller University, 14. Brown University, 15. University of Colorado Anschutz Medical Campus, 16. Swiss Federal Institute of Technology, Lausanne, 17. New York University, 18. La Jolla Institute, 19. IMGT, Univ Montpellier, 20. St. Jude Children’s Research Hospital, 21. Alternative Energies and Atomic Energy Commission, Grenoble, 22. Massachusetts Institute of Technology

## Methods

### Animals

All mice used in the generation of the immgenT dataset are referenced in Extended Data Table 1, including sex, age, and genetic background. With rare exceptions, experiments were performed using C57BL/6 (B6) mice, most of which were sourced from The Jackson Laboratory. Experimental conditions, including infection models, immunization strategies, and tissue processing protocols, are detailed in Extended Data Table 1, the GEO GSE297097 dataset, and the immgenT websiteMice for experiments were, with a few exception all B6J, sex-balanced. age-matched, littermates when possible.All mice were initially bred and maintained under specific pathogen–free conditions in accordance with the animal care and use regulations of each institution where the experiments were performed. All experimental procedures were approved by the Institutional Animal Care and Use Committee of each institution.

## Single-cell RNAseq, TCRseq, and CITEseq staining - Sample Preparation

The following paragraphs describe the overall strategy for all immgenT experiments. Sample-specific variations are described in **Extended Data Table 1**, with specific columns referenced throughout the text. A step-by-step protocol is available on the immgenT website (https://www.immgen.org/ImmGenT/immgenT.SOP.pdf).

Each dataset in the project was assigned an identifier, IGT1-96, as described in Extended Data Table 1, Column *IGT.* This IGT code is used throughout to track individual datasets, and starts the unique cell identifier (IGT.cellID) used as the linking key. Hashtag were used to tag individual samples. IGT.HT (e.g., I20H5) uniquely identifies samples.

Each experiment included a spleen standard, consisting of separately labelled splenocytes from a 6-8 week JAX B6 mouse, male or female, to facilitate batch correction and comparison. These are flagged in Extended Data Table 1 Column *spleen_standard*.

IGT code is used throughout to track individual datasets, and starts with the unique cell identifier (IGT.cellID) used as the linking key. Hashtags were used to tag individual samples. IGT.HT code is used throughout to track individual samples.

### Logistics

Experiments were conducted across multiple laboratories in the United States. For each experiment, an immgenT research assistant coordinated sample collection and performed encapsulation at the host institution. Participating laboratories carried out mouse treatments (e.g., infection or immunization) and tissue processing, with assistance for hashtagging, CITEseq staining, and encapsulation. Each experiment typically included ∼10 pooled samples and a standardized spleen control for batch-effect assessment. Library construction and sequencing were centralized at the Broad Institute.

### Flow cytometry and hashtag antibody staining

Single-cell suspensions from tissues were isolated by mechanical dissociation or enzymatic digestion and stained simultaneously with flow cytometry antibodies and different TotalSeq-C Anti-Mouse Hashtags^101^ (TotalSeq-C Anti-Mouse Hashtags 1-24 Biolegend #155861-155879, #113931-113939), in staining buffer (phenol red-free DMEM, 2% FCS and 10mM HEPES) for 20min at 4°C in the dark.

For the spleen standards, splenocyte suspensions were stained under identical conditions and included an anti-CD45 antibody conjugated to a unique fluorophore, thereby allowing them to be differentiated from the other samples, even when subsequently pooled during sorting. Each sample was labeled with a unique hashtag, enabling the downstream assignment of each individual cell (10x cell barcode) to their respective source samples, as described in Extended Data Table 1 column *HT*. Most samples represent a single mouse and tissue. In some cases, low cell numbers required pooling multiple mice per sample, as detailed in Extended Data Table 1 column *pool_of_mice*.

### Cell sorting

Each sample and spleen standard was sorted individually by flow cytometry (e.g., FACSAria) and pooled in a single collection tube at 4 °C, to a combined total of up to 500,000 T cells. DAPI was added just before the sort to exclude dead cells. In most samples, live DAPI-CD3+ T cells were sorted. Specific populations were sorted as described in Extended Data Table 1 column *gating_strategy*. Spleen standards were sorted as DAPI-CD45+ or DAPI-CD3+.

### ImmGenT TotalSeq-C custom mouse panel staining (CITEseq)

A lyophilized tube of ImmGenT TotalSeq-C custom mouse panel, containing 128 antibodies (Biolegend Part no. 900004815, panel information detailed in Extended Data Table 2) was equilibrated to room temperature for 5min; centrifugated for 30sec at 10,000g; resuspended with 24uL of staining buffer (described above) and 1uL of FcBlock (Bio X Cell #BE0307); and incubated at room temperature for 30min. The tube was then centrifugated for 10min at 14,000g to pellet aggregated antibodies. 25uL of the mixture was aspirated and used to stain the sorted cells.

The CITEseq staining protocol requires 500,000 cells; when necessary, unstained splenocytes were spiked into the collection tube to reach this target. The cells were resuspended in 25 μL of staining buffer, to which 25 μL of aspirated CITEseq mixture was subsequently added. The cells were stained for 20min at 4°C in the dark, then washed 4 times with 1mL of staining buffer to remove unbound antibodies.

### Second sort

Cells were subsequently sorted a second time to remove unbound antibodies. DAPI and the distinct CD45-conjugated fluorophores were used to sort up to 55,000 live T cells and 5,000 live spleen standard cells into a single collection tube. The cells were resuspended in 38.7 μL of staining buffer, then immediately taken for encapsulation.

### Single-cell RNAseq, TCRseq and CITEseq-Sequencing - Library Preparation

#### Cell encapsulation and cDNA library

Single-cell RNA sequencing was performed using the 10x Genomics 5’ v2 platform with Feature Barcoding for Cell Surface Protein and Immune Receptor Mapping, adhering to the manufacturer’s guidelines (CG000330). After cell encapsulation with the Chromium Controller, reverse transcription was performed on the emulsion, and the resulting cDNA was amplified via PCR. The amplified cDNA library was separated on the basis of fragment size using SPRISelect beads (Beckman Coulter, B23317). Larger, transcript-derived fragments were preserved for TCR and Gene Expression library construction, while smaller, TotalSeq-C-derived cDNA fragments were used for Feature Barcode library construction.

#### Gene expression (GEX) library construction

After enzymatic fragmentation and size selection of the cDNA, the library was ligated to an Illumina R2 sequence and indexed with a unique Dual Index TT Set A (10x Genomics PN-3000431) index sequence.

#### TCR library construction

The alpha beta TCR transcripts were amplified from the transcript-derived cDNA using a nested PCR utilizing primers provided in the 10x Genomics Single Cell Mouse TCR Amplification Kit (PN-1000254). The enriched TCR transcripts were fragmented and size-selected, ligated to an Illumina R2 sequence, and indexed with a unique Dual Index TT Set A (10x Genomics PN-3000431) index sequence.

#### TotalSeq-C library construction

Totalseq-C-derived cDNA (CITESeq panel, Hashtags) was processed into the Feature Barcode libraries following the manufacturer’s protocol. The library was indexed with a unique Dual Index TN Set A (10x PN-3000510).

#### Quality control

Fragment sizes for both cDNA fractions, intermediate TCR amplification pools, and finished libraries were evaluated using the Agilent Bioanalyzer 2100 High Sensitivity DNA assay, while concentrations were quantified with a Qubit dsDNA HS Assay kit on a Qubit 4.0 Fluorometer.

#### Sequencing

Libraries were sequenced on either an Illumina Novaseq6000 or, once available, Novaseq X sequencer. On the Novaseq6000, the three library types for each 10x lane were pooled based on molarity in the following proportions: 47.5% RNA, 47.5% Feature Barcode, and 5% TCR. On the Novaseq X, accounting for the higher read yield and preferential sequencing of shorter fragments, the three library types for each 10x lane were instead pooled in the following proportions: 60% RNA, 25% Feature Barcode, and 15% TCR. In all cases, the pooled libraries were sequenced using 10x Genomics-recommended specifications: 26 cycles for Read 1, 90 cycles for Read 2, 10 cycles for Index 1, and 10 cycles for Index 2.

### Single-cell RNAseq and CITEseq - Data processing

#### Count matrices

Gene and TotalSeq-C antibody (surface protein panel and hashtags) counts were obtained by aligning reads to the mm10 (GRCm38) mouse genome using the M25 (GRCm38.p6) Gencode annotation and the DNA barcodes for the TotalSeq-C panel (in Extended Data Table 1). Alignment was performed with CellRanger software (v7.1.0, 10x Genomics) using default parameters. Cells were identified and separated from droplets with high RNA and TotalSeq-C counts by determining inflection points on the total count curve, using the barcodeRanks function from the DropletUtils package^102^.

#### Sample demultiplexing

Sample demultiplexing was performed using hashtag counts and the HTODemux function from the Seurat package (Seurat v5^103,104^). Doublets (droplets containing two hashtags) were excluded, and cells were assigned to the hashtag with the highest signal, provided it had at least 10 counts and was more than double the signal of the second most abundant hashtag. Hashtag count data were also visualized using t-SNE to ensure clear separation of clusters corresponding to each hashtag. All single cells from the gene count matrix were uniquely matched to a single hashtag, thereby linking them unambiguously to their original sample.

#### Quality control (QC)

Cells were excluded if they met any of the following standard criteria: fewer than 500 total RNA counts; greater than 10% of counts mapping to mitochondrial genes (indicative of low-quality or dying cells); fewer than 500 TotalSeq-C counts; or positivity for two isotype control antibodies, consistent with non-specific TotalSeq-C binding. Non–T cells were further removed based on gene expression signatures, including myeloid/natural phagocyte (MNP), B cell, and innate lymphoid cell (ILC) programs, together with absence of a T cell gene signature. Signature scores were computed using AddModuleScore_UCell^105^. Additional details are provided in Extended Data Note 1 and 2. In experiments where CITEseq data did not meet these quality control criteria, protein measurements were excluded from downstream analyses; such experiments are indicated in Extended Data Table 1 (column *cite_seq*). Further details are described in Extended Data Note 2.

### Data integration, dimensionality reduction, and clustering and annotation

#### Data integration

immgenT datasets (IGT1–96) were integrated using the scVI/totalVI model (v1.2.0)^25^, with the 10x Genomics lane (IGT) specified as a batch covariate. The model was trained with default parameters using all detected genes and proteins, with a 30-dimensional latent space.

#### Dimensionality reduction

Dimensionality reduction of the 30-dimensional totalVI latent space for visualization was performed using Minimum Distortion Embedding (MDE) implemented in the pyMDE^33^ library via the pymde.preserve_neighbors() function with default parameters. In contrast to UMAP, which is stochastic and graph-based and primarily optimized for visualization, pyMDE preserves both local neighborhood structure and global geometry with minimal distortion. After computing the initial embedding on the full immgenT dataset (IGT1–96), the coordinates were reused to anchor immgenT cells as a reference, enabling projection of new datasets without altering the original embedding. This enabled the construction of shared T-cell reference embeddings (both all-T and lineage-specific) for comparative analyses.

#### Lineage annotation (“level 1”)

Cell clustering was initially performed to assign cells to major immune lineages. Primary lineage annotation (CD4, CD8, Treg, γδ T, Tz, DN, and DP) was based on combined RNA and protein expression of canonical markers, including *Cd3e*, *Trbc1*, *Trbc2*, *Cd4*, *Cd8a*, *Cd8b1*, *Foxp3*, *Zbtb16*, *Trgc1*, and *Trdc*, as well as surface proteins CD3, TCRB, THY1.2, CD4, CD8A, CD8B, and TCRGD. Clustering was performed in the totalVI latent space using Louvain’s algorithm with Seurat’s FindClusters() across a range of resolutions (0.1–4) to ensure that clusters corresponded to single lineages. Initial annotations were assigned at the cluster level and subsequently validated at single-cell resolution using CITEseq data. Approximately 1% of cells showed discordant marker expression (e.g., CD4 expression within CD8 clusters) and were manually corrected. iNKT and MAIT cells identified by TCR sequencing were in Tz clusters, with ∼1% manually reassigned.

#### Fine-grained clustering (“level 2”)

For each major lineage, cells were reclustered independently using the same overall totalVI latent space (as the initial model was trained jointly on all genes, proteins, and cells, rerunning totalVI within individual lineages did not improve granularity). Clustering was performed by evaluating Louvain clustering solutions across multiple resolutions (0.5–4), with silhouette analysis used to minimize over-and under-clustering (typically optimal at resolutions 0.75–1). Large clusters that were oversplit were merged using lower-resolution solutions, and smaller populations better resolved at higher resolution were split. Decisions to merge or split clusters were made collectively by the immgenT consortium during a series of four workshops, based on consistency across samples, coherence of gene-expression signatures, and protein expression profiles. In total, 1,710 of 719,580 cells (0.02%) scattered in small clusters could not be confidently annotated and were annotated as “unclear”. The analysis excluded them. Of note, cluster numbers are not contiguous, reflecting their historical assignment during iterative analyses.

#### level2_group annotation

Resting versus activated states were defined based on expression of *Sell* (CD62L) and *Cd44* (CD44). Proliferating cells were identified based on *Mki67* expression and cell-cycle scoring (see below). “Miniverse” cells formed separate clusters labeled wM.

ImmgenT integration and annotation are available on https://www.immgen.org/ImmGenT/. (run_totalvi_v2.py)

#### UMAP and clustering of individual IGT datasets

UMAP and clustering were also performed on individual IGT datasets, generating dataset-specific UMAPs used in the Rosetta2 databrowser and in some immgenT manuscripts. These used the standard Seurat pipeline: NormalizeData(normalization.method = “LogNormalize”, scale.factor = 1e4) %>% FindVariableFeatures(selection.method = “vst”, nfeatures = 2000) %>% ScaleData(features = VariableFeatures(.)) %>% RunPCA(features = VariableFeatures(.), npcs = 50) %>% FindNeighbors(dims = 1:30) %>% FindClusters(resolution = 1).

### Updating MDE embeddings with new datasets

Existing Minimum Distortion Embedding (MDE) visualizations were updated using an anchored optimization strategy implemented in *pymde 0.2.0*^33^ (default parameters). Reference cells with a precomputed two-dimensional embedding were fixed using pymde.Anchored(), while new cells were added by optimizing a neighbor-preserving objective defined with pymde.preserve_neighbors(). Nearest-neighbor relationships were computed in the shared latent space, and anchoring constraints ensured that reference cell coordinates remained unchanged during optimization. This approach preserves the global geometry of the original embedding while allowing consistent placement of newly integrated cells, enabling incremental extension of the MDE without recomputing the full layout. (run_mde_incremental.py)

### Cluster Robustness Analysis

#### Cluster Separability Score (CSS)

To quantify transcriptional distinctiveness at the cluster level, we computed a separability score for each cluster within each dataset. Cells were embedded in PCA space for each IGT (RNA or Protein space), and cluster centroids were defined as the mean coordinates of all cells in a cluster. Separability was defined as the difference between the mean distance of a cluster centroid to all other cluster centroids and the mean intra-cluster distance of cells to their own centroid. This metric captures global transcriptional separation while being robust to cluster size and local neighborhood structure.

For each cluster CCC within a dataset, we defined a normalized cluster separability score as S(C) = \frac{B(C) - A(C)}{\max(A(C), B(C))},S(C)=max(A(C),B(C))B(C)−A(C),

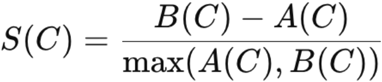

where A(C)denotes the mean intra-cluster distance of cells to their cluster centroid, and B(C) denotes the mean distance between the cluster centroid and all other cluster centroids in PCA space. This metric is bounded between −1 and 1, with positive values indicating that clusters are more transcriptionally separated from other clusters than they are internally dispersed.

#### Intra-cluster gene expression consistency

To quantify robustness within each immgenT cluster we assessed the consistency of gene-expression behavior across experiments (IGT). For each cluster, the algorithm selects two IGTs and derives a weight vector of length equal to the number of genes, where each weight is inversely proportional to the Wasserstein distance between the gene expression distributions of the two. We then compute the cosine similarity across all pairs. The robustness is plotted as the distribution of all cosine similarities. A null distribution is generated by randomly reassigning cell clusters and running the same algorithm (n=1,000 times). This metric directly evaluates uniformity and consistency of gene expression within a cluster, independent of other cell types.

### Differential Gene Expression

*Limma*. Differential gene expression was performed using a pseudobulk linear modeling framework based on limma^106,107^ and edgeR. Briefly, raw RNA counts from the Seurat object were aggregated by cell cluster (annotation_level2) and experiment (IGTHT), and normalized using trimmed mean of M-values (TMM). A design matrix encoding cluster–experiment combinations was constructed, and gene-wise linear models were fit to the normalized expression matrix using limma. Differential expression was assessed using empirical Bayes moderation of variance estimates, and statistical significance was determined with Benjamini–Hochberg correction for multiple testing.

(limma_wrapper_template.sh limma_make_tmm_template.R limma_fit_template.R limma_contrasts_template.R)

*FlashierDGE*. Efficient differential expression was performed in the flashier EBMF semi-NMF framework (GP model, see below) by leveraging the learned gene programs (factors) and cell loadings from a 200-factor model fit to log-normalized expression. For a given comparison, mean factor loadings were computed separately for group 1 and group 2, and the difference in mean loadings (Δloadings) provided differential gene-program activity. Gene-level differential expression was then reconstructed directly from the factorization as F×Δloadings, yielding per-gene log fold changes. The same framework also returned average expression estimates for both programs and genes from the group-wise mean loadings and reconstructed group means. The function is available in ZemmourLib package (FlashierDGE)

### Gene signature scoring

Gene signature scores were computed using Seurat’s AddModuleScore() function. Tissue-resident memory T cell signature was derived from Milner *et al.*^89^ Core regulatory T cell (Treg) identity was assessed using a curated gene set from Zemmour *et al.*^90^: *Foxp3, Ikzf2, Il2ra, Ctla4, Il2rb, Capg, Hopx, Tnfrsf4, Tnfrsf18, Tnfrsf9,* and *Izumo1r*. Stress-related signatures were obtained from the following sources: dissociation signatures from van den Brink et al.^39^; an immediate-early response signature from Bahrami et al.^108^ (*Nr4a1, Egr1, Egr2, Jun, Junb, Fos,* and *Fosb*); interferon-stimulated gene (ISG) signatures from ImmGen^109^, as well as GP12 and GP16 from the immgenT gene-program analysis; mitochondrial genes (all genes beginning with “Mt-”); an integrated stress response (ISR)/chronic ATF4-signaling signature from Alicea Pauneto et al.^110^ (*Atf4, Ddit3, Ero1a,* and *Ppp1r15a*) and GO:0140467; an ER-stress signature from Chow et al.^111^, together with response to endoplasmic reticulum stress (GO:0034976) and endoplasmic reticulum unfolded protein response (GO:0030968); and a hypoxia signature from Bono et al.^112^, together with response to hypoxia (GO:0001666).

### Cell-cycle scoring

Cell-cycle activity was inferred using phase-specific gene sets from Dominguez *et al.*^113^. Following log-normalization, S-phase (G1–S) and G2/M scores were computed per cell using Seurat’s CellCycleScoring() function with mouse orthologs, enabling identification of proliferating cell populations.

### CITEseq marker combination for immgenT clusters

For each lineage, COMETSC^114^ was applied with default parameters to normalized protein expression matrices together with MDE coordinates and cluster annotations, allowing up to two-gene combinations (-K 2) to identify combinatorial markers. Following manual inspection, extended marker combinations (not limited to two genes) were evaluated for sensitivity (Se), specificity (Sp), positive predictive value (PPV), and negative predictive value (NPV), and were subsequently validated by flow cytometry in each companion manuscript (immgenT-Treg^46^, immgenT-CD8^45^, immgenT-CD4^47^, immgenT-Tz)

### Immgen T reference-based integration (T-RBI)

T-RBI integrates and maps external scRNA-seq datasets onto the immgenT atlas using Scanpy, scVI/scANVI^50^, and pyMDE^33^. Reference (immgenT) and query datasets are prepared for integration with AnnData (v0.11.4), Scanpy (v1.11.2), and pandas (v2.3.0). Only transcriptomic data are used (no other modalities, such as CITEseq). T-cell and γδ T-cell (gdT) signature scores are calculated separately with Scanpy’s *score_genes* function. Query cells with a T-cell score <0 are excluded, and cells with a gdT score >0 are assigned to the gdT lineage.

### Initial scVI models

gdT and αβ T-cell (abT) populations are then separated into independent AnnData objects and modeled separately using scVI (v1.3.1.post1). For each population, data are prepared using *prepare_anndata* (*batch_key* = “IGT|query dataset”, *categorical_covariate_keys* = [“IGTHT|query samples”]), followed by *scvi.model.SCVI*(*n_latent* = 30, *n_layers* = 2) and *model.train()*. The resulting scVI latent representations are retained for subsequent visualization with pyMDE (v0.2.1) and the trained scVI models are used to initialize scANVI models for annotation.

### Cell annotation

For abT cells, scANVI is applied sequentially for lineage (level 1) and sub-lineage (level 2) prediction using *scvi.model.SCANVI.from_scvi_model* with query cells initially annotated as “*unknown*” in *labels_key* = “level1_abT” or “level2_abT”, respectively, and *model.train(max_epochs = 25)*. Because gdT lineage identity has already been assigned using the gdT signature score, scANVI is applied to gdT cells only for sub-lineage (level 2) prediction using *labels_key* = “level2_gdT”. We use *model.predict* from each scANVI model to get our predicted annotations and their confidence score. Using this prediction score, we assign those cells with a score < 0.85 as “not classified”. We then fine-tune the annotation. We use only the query cells and run the above scVI/scANVI models to annotate “not classified” cells (lineage and sub-lineage for abT cells; sub-lineage for gdT cells) based on the already labeled query cells. Fine-tuning is iterated until one of the following conditions is met: < 1% “not classified” or < 33% change between iterations. We output the annotations and confidence score before and after fine-tuning. We then calculate the discovery score for the remaining “not classified” cells, as described below.

### ImmgenT embeddings

To embed the query cells into our existing immgenT MDEs (allT MDE and lineage-specific MDE), we use pyMDE, anchoring reference cells *(embedding_dim=2, constraint=anchor_constraint, repulsive_fraction=0.7, n_neighbors=15, verbose=True, device=’cpu, eps=1e-6).* For the all-T MDE embedding, we use the initial scVI latent spaces computed separately for αβ and γδ T cells. For lineage-specific MDEs, we use the scANVI level 2 latent representations for each lineage. Lineage-specific embeddings are not generated for lineages represented by fewer than five cells.

Output files include level 1 and level 2 MDE coordinates, cell annotations with confidence scores before and after fine-tuning, and discovery scores. A PDF summary of the results is also provided, displaying the all-T MDE, lineage-specific MDEs, and corresponding annotations. The distribution of confidence scores after tuning is shown as a density plot. Unannotated cells (score < 0.85 after fine-tuning) are visualized in a dataset-specific UMAP to assess whether they form a distinct cluster. Discovery scores are shown on the all-T MDE. Options are available to map to specific lineages rather than the whole dataset.

### k-nearest neighbor discovery score

To quantify whether unannotated cells occupy regions of the embedding space not represented in the annotated reference, we defined a local k-nearest neighbor (kNN) distance ratio score. For each dataset, principal component analysis (PCA) was performed independently, and all distances were computed in the resulting dataset-specific 10-dimensional PCA space using Euclidean distance.

For each query cell *qi*, we computed (i) the mean Euclidean distance to its k=10 nearest neighbors among the annotated cells and (ii) the mean Euclidean distance to its k=10 nearest neighbors among the unannotated cells, excluding the cell itself. The novelty score was defined as the ratio of these two quantities:

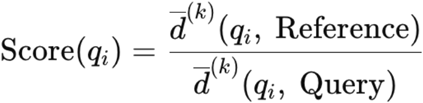

where *d^-(k)^*(*qi,S)* denotes the mean distance from *qi* to its *k* nearest neighbors in set SSS. Scores greater than 1 indicate that an unannotated cell is locally closer to other unannotated cells than to annotated cells, consistent with occupancy of a region of the embedding space under-represented or new in the reference. Conversely, scores near or below 1 indicate that unannotated cells are embedded within previously annotated regions.

### T-RBI cluster ablation tests

To test whether T-RBI can recognize T-cell states that are absent from the reference, we performed leave-one-cluster-out ablation analyses using the CD4 lineage. For each indicated CD4 cluster, all cells belonging to that cluster were removed from the immgenT reference and treated as an independent query dataset. The remaining CD4 cells, with their original cluster annotations, constituted the ablated reference. The same query cells were also mapped against the complete, non-ablated CD4 reference as a control. For each ablation, query cells were processed using the standard T-RBI pipeline described above, including scVI/scANVI integration, level 2 cluster prediction with and without fine tuning, discovery calculation based on fine-tuned annotations, and projection onto the anchored CD4 MDE.

We quantified the outcome of each ablation by recording the fraction of query cells assigned to each remaining CD4 cluster or left unclassified, and the corresponding annotation confidence scores before and after fine-tuning. Discovery scores were calculated for the ablated query cells using the same k-nearest-neighbor procedure described above and compared with scores obtained when the same cells were mapped against the complete reference. A discovery score >1.1 was used to flag cells preferentially associated with other query cells rather than with annotated reference cells, consistent with a state under-represented or absent from the reference.

### Gene expression sensitivity and specificity across clusters - AUC analysis

To identify transcription factors and effector molecules that are both sensitive and specific across clusters, we computed an area-under-the-curve (AUC)–based enrichment score from pseudobulk expression profiles. Log-normalized pseudobulk expression values (log1p CP10K) were obtained using Seurat’s AggregateExpression() function. For each gene, expression values were normalized by the gene’s total expression across all clusters, such that the normalized expression profile summed to one.

Clusters were then ranked in descending order of normalized expression for each gene, and the cumulative sum of normalized expression was computed along this ranked list. The AUC score was defined as the mean of the cumulative expression values across ranks. This score quantifies how selectively a gene is enriched in a limited number of clusters versus broadly distributed across many clusters. Ranging from 0 to 1, genes whose expression is concentrated in a small number of clusters achieve higher AUC scores, whereas genes with diffuse or uniform expression across clusters yield lower AUC values. Genes with expression exceeding a minimal threshold (0.5 log1p CP10K) in at least one cluster were retained for analysis.

This analysis was performed either across all clusters or within clusters stratified by lineage, and was applied to both transcription factors and effector molecules.

The transcription factor list was obtained from Gene Ontology^115,116^ GO:0003700 (org.Mm.eg.db package version 3.20.0), with the additional inclusion of *Tox*, *Tox2*, *Tox3*, and *Tox4*. Effector molecules were curated from Gene Ontology GO:0005615.

### Gene set enrichment analysis

Gene set enrichment analysis (GSEA) was performed using ranked gene-level statistics and Gene Ontology (GO) gene sets obtained from MSigDB via the msigdbr R package^117^. GO collections from MSigDB category C5 were analyzed separately for Biological Process (GO:BP), Molecular Function (GO:MF), and Cellular Component (GO:CC). Enrichment was computed using fgseaMultilevel with scoreType = “pos” and minSize = 5. Enrichment results were summarized using the normalized enrichment score (NES) and multiple-testing–adjusted P values (FDR; padj), with significance defined as FDR < 0.05.

### Gene Programs - Empirical Bayes matrix factorization of gene expression

To identify latent transcriptional programs, we applied empirical Bayes matrix factorization (EBMF) with a semi–non-negative matrix factorization (semi-NMF) structure to log-normalized single-cell RNAseq expression using the flashier framework^85,118^. EBMF decomposes an expression matrix into a sum of low-rank components while learning sparsity, shrinkage, and noise parameters directly from the data via empirical Bayes, enabling robust identification of interpretable gene programs without requiring a priori selection of factor number or regularization strength. In the semi-NMF formulation used here, gene loadings are constrained to be non-negative, facilitating biological interpretability of gene programs, while cell loadings are unconstrained.

We used the standard pipeline of *flashier* for single-cell RNAseq. Gene expression was log-normalized using Seurat’s LogNormalize() with a scale factor set to the mean total UMI count per cell (mean(nCount_RNA)), yielding log1p-transformed expression values. Genes were filtered prior to model fitting to reduce technical and clonotype-driven variation: TCR variable/constant genes (e.g., Trbv/Trbj/Trbc, Trav/Traj/Trac, Trgv/Trgc, Trdv/Trdc), mitochondrial genes (^mt-), ribosomal genes (including Rpl/Rps/Mrpl/Mrps/Rsl), predicted/RIKEN/pseudogene-like features (including Gm genes, genes ending in Rik, and-ps pseudogenes), and genes not detected in any cell (row sum of counts = 0) were excluded.

Variance regularization parameters were set using a reference Poisson sampling procedure: a Poisson distribution with rate 1/n (where n is the number of cells) was sampled to estimate the standard deviation of log(x+1) and used as the residual noise scale parameter (S). EBMF components were fit using flash() (flashier^118^) with a mixture of empirical Bayes priors (ebnm_point_exponential and ebnm_point_laplace) and variance model var_type = 2, with a maximum of 200 greedily added factors (greedy_Kmax = 200). Backfitting was enabled to refine factor estimates.

Latent factor scores were analyzed using linear modeling with the limma framework. A design matrix was constructed, including cell state identity (annotation_level2) as the primary variable of interest and experimental replicate (IGT) as a covariate to control for confounding effects. For each factor, a linear model was fit using lmFit, followed by empirical Bayes moderation with eBayes (trend = TRUE, robust = TRUE). For each cell state, contrasts were defined to compare that cluster against the average of all other clusters (one-vs-all). Differential factor loadings were assessed using moderated t-statistics, with p-values adjusted for multiple testing using the Benjamini–Hochberg procedure. Effect sizes (contrast coefficients) and adjusted p-values were visualized as heatmaps, with significant associations (FDR < 0.05) annotated.

### Plotting

Plots were generated in R using ggplot2^119^, S-Plus, Seurat^104^, or the ZemmourLib R package (https://github.com/dzemmour/ZemmourLib, v0.1.3). Heatmaps were generated using pheatmap (v1.0.13) or Morpheus (Broad Institute)

## Code availability

Code is available at the following repositories: https://github.com/immgen/immgenT_Project/tree/main/Data_processing_pipeline, https://github.com/immgen/immgenT_Project/tree/main/Cosmology_paper, https://github.com/immgen/immgenT_Project/tree/main/GP_paper.

## Data availability

All raw and processed sequencing data generated in this study are available through the Gene Expression Omnibus (GEO) under accession GSE297097.

Studies mapped using T-RBI listed in **Extended Data Table 5**

Additional resources are described in detail in **Extended Data Note 2**.

## immgenT resource: public data browsers

Beyond the data and analyses reported in this and accompanying articles, immgenT’s goal as a public resource is served by a set of data browsers which enable multi-faceted exploration of the data. They are accessible through the immgenT portal (https://www.immgen.org/ImmGenT/) and include: **Rosetta2 (**https://rosetta.immgen.org/) is an interactive single-cell viewer that connects RNA-seq and protein (CITE-seq) data. Cell populations can be visualized on parallel RNA and protein UMAPs, coloring reciprocally RNA-or protein-based clusters. The expression of individual genes, or of user defined gene signatures can be mapped on these representations. A cell gating interface modeled after flow cytometry serially generates 2D CITE-seq plots, on which cells can be “gated”, for projection onto UMAPs or computation of differential expression. Rosetta2 returns differentially expressed genes and proteins (as heatmaps or tables). Rosetta2 can be used to explore individual datasets or, or integrated cells grouped by lineage.

**TCR Browser** (https://rstats.immgen.org/tcrbrowser/) is a searchable interface for TCR sequences across the immgenT data. TCR repertoires can be browsed within immgenT datasets, searched for TCRs that use specific V, J or CDR3 elements, or for similarity to a user provided TCR.

**T-RBI portal** (Reference Based Integration) (https://rstats.immgen.org/immgenT_integration_analysis/) allows AI-driven mapping of external single-cel RNAseq datasets into the immgenT framework. Users upload their own data and metadata files (standard sc formats), which are then mapped and annotated within the immgenT framework. After an overnight compute, the tool returns annotation tables with integration statistics and annotation of every cell in the immgenT cluster framework, and x/y coordinates in the reference MDE space.

**Skyline browser** (https://rstats.immgen.org/Skyline/skyline.html?datagroup=immgenT%20Pseudobulk&) represents, as a classic bar histogram, a pseudo-bulked overview of expression profiles of a given gene across all or selected immgenT cell clusters

## Extended Data

### Extended Data Note 1 - Quality Control

#### Quality control

Each experiment underwent stringent quality control to ensure robust and comparable measurements across samples. Cells were retained according to standard single-cell RNAseq criteria, including adequate gene and UMI coverage, low mitochondrial RNA content, and exclusion of doublets (**Methods, Extended Data Fig. 1, Extended Data Table 3**).

A subset of cells consistently clustered into a distinct state, which we termed the miniverse (wM clusters). While multiple analyses argue against a technical origin, we cannot fully exclude the possibility that this state reflects, in part, technical or biological edge cases, and it is therefore interpreted cautiously.

Mature double-positive (DP) T cells outside the thymus comprised a small fraction of the dataset. Although some of these cells likely represent bona fide peripheral T cell states, and cells clearly identified as doublets were removed (**Methods**), we cannot completely rule out residual technical artifacts within this population.

#### CITEseq performance analysis

For CITEseq^29,30^, we leveraged isotype control counts included in the antibody panel to identify and filter cells with elevated non-specific background signal. In four experiments, CITEseq staining failed globally, as indicated by comparable antibody-derived tag (ADT) counts in cell-containing and empty droplets; these experiments were excluded from downstream protein-level analyses.

Evaluating the performance of individual antibodies in the CITEseq panel was not trivial, since the signal reflects not only the quality of the reagent but also the actual protein expression levels (which are unknown). The relationship to mRNA intensity in the same cells can serve as a guide, but is confounded by translational and post-translational regulation. We used two metrics: (i) the dynamic range of protein signal and (ii) comparison of surface protein abundance with transcript expression across the dataset, computing RNA–protein correlations for each antibody (**Extended Data Fig. 2a-d, Extended Data Table 4**). As illustrated (**Extended Data Fig. 2a,c**), antibodies yielded different profiles. Some high-performing antibodies (e.g., CD4, CD8β, Ly49A exhibited both strong RNA–protein correlation and broad dynamic range, producing expression patterns closely resembling flow cytometry (see also the thymus staining profiles in the accompanying immgenT-TCR manuscript^120^) and enabled clear separation of negative and positive populations or graded levels of expression that tracked with RNA levels in the cell clusters. Others showed intermediate performance with a more limited dynamic range (e.g., CD127/IL7-RA), or with detectable protein expression primarily at higher transcript levels (**Extended Data Fig. 2b)**. A third group showed poor performance (**Extended Data Fig. 2c)**, with little or no detectable protein signal despite transcript expression, and were excluded from protein-based interpretations. Most chemokine receptors were not robustly detected, perhaps predictably because many antibodies to chemokine receptors require specific staining conditions (e.g., staining at 37°C for an extended period of time), not applicable for a broad panel. The plot of (**Extended Data Fig. 2d)** summarizes these observations and should provide a valuable guide to interpretation and for future experiments.

Beyond intrinsic antibody performance, we identified experimental factors that influenced the CITEseq signal. Specifically, competition between antibodies targeting the same epitope during staining and sorting reduced sensitivity, particularly for CD3, CD3e, and CD45, which were extensively used for cell enrichment across experiments. In contrast, enzymatic tissue digestion had a minimal impact on epitope integrity for most markers (data not shown).

Overall, the CITEseq panel performed robustly across most experiments and enabled true single-cell protein quantification that closely parallels flow cytometry. Unlike transcriptomic measurements, CITEseq does not suffer from dropout, and for high-performing antibodies, protein expression distributions permitted conventional gating strategies, effectively bridging high-dimensional single-cell sequencing with classical immunological analyses. Together, these results demonstrate that immgenT provides a rigorously quality-controlled, protein-and transcriptome-resolved resource in which each experiment can be analyzed independently or in aggregate.

#### Extended Data Note 2 – immgenT public web resources

Beyond the datasets generated and the papers that describe their analysis, the immgenT program has deployed four new online tools to support the public resource aspect. These are accessible through the immgenT portal (https://www.immgen.org/ImmGenT/). The reference cell-state clusters and their representation on Minimal Distortion Embedding (MDE) maps generated by immgenT provide a common reference for these tools, allowing a common language to describe T cells that applies across many locations and conditions.

**Rosetta2 (**https://rosetta.immgen.org/)

Rosetta2 is a multi-faceted visualization and exploration tool for single-cell data that enables users to explore cell clusters and gene expression in immgenT datasets or in integrated datasets (either “allT”, or lineage-specific projections). Rosetta2 uniquely enables side-by-side exploration of RNA and protein data, and representations derived from flow cytometry to select cells based on protein marker data.

The user first chooses datasets from individual immgenT experiments (IGTs), or integrated lineage-specific data (CD4, CD8, Treg, etc., or allT). Once a dataset is chosen, four pages are available to explore the data: Dataset Overview, Flow-like Genomics, Subset Comparison, and Integrated Data.

1. The dataset overview acts as a landing page and compares cell clusters in the mRNA and protein space, filtering or coloring cells according to sample origin or cell state. The expression of individual genes or of user-defined gene signatures can be shown on the MDE or UMAP representations. Markers of specific populations (clusters or user-defined regions of interest) can be displayed as heatmaps (top 15 upregulated and downregulated genes, for RNA or protein separately).
2. Flow-like Genomics treats the CITEseq data as if from a Flow Cytometry experiment, with the functionalities common to flow cytometry software: selection of a group of cells on serial 2D plots, “gating” to filter or highlight those cells on subsequent panels, and projecting the selected cells onto the MDE maps. The gating path can be related back to the UMAPs in both RNA and Protein space, filtering cells based on the Boolean combination of gates.
3. Subset Comparison allows a differential gene expression analysis between two selected cell subsets of cells, reported as a heatmap.
4. Group Integration allows the user to project cells from the current dataset onto the full “all-T” integrated reference (MDE plot). Expression of selected genes or gene signatures can be shown on the two spaces.

**T-RBI Data Integration (**https://rstats.immgen.org/immgenT_integration_analysis/) The immgenT T-RBI (“Reference-Based Integration”) page provides a platform for researchers to integrate their own scRNAseq T cell datasets into the immgenT reference framework (either the complete reference or specific lineages, e.g., CD4 and CD8). T-RBI returns two complementary outputs: (i) annotations into immgenT reference clusters with associated confidence and discovery scores for each query cell; (ii) coordinates of query cells within the reference immgenT MDE graphs.

The integration pipeline utilizes scVI/scanVI^24,50^ (from scVerse) for data harmonization and pyMDE^33^ to project query cells into the reference MDE space. As of this writing, the tool has been tested mainly on single-cell RNA-seq data generated on the 10X Genomics platform (3’, 5’), but consistent results have been obtained with other platforms (e.g., Flex, Parse). To manage computational resources, submissions are currently limited to one job per 24 hours per user, with a maximum capacity of 25,000 cells per job. Larger datasets could be partitioned into multiple submissions.

Users are asked to upload standardized inputs, including a raw counts matrix, feature names, and cell barcodes (generated natively by CellRanger/10x Genomics), and a cell-level sample metadata file. Users also enter the lineages of those cells (if known). Following an overnight compute period, users receive by email a comprehensive results package, including: 1. A PDF report visualizing the success of the integration, distribution of confidence scores for cells that have been assigned, or a “discovery score” for those cells that T-RBI cannot unambiguously assign. 2. An output file containing the user’s original cell IDs with immgenT lineage and cluster memberships, the confidence score and discovery score. 3. Coordinates of cell in both the all-T cell and lineage-specific immgenT MDE plots.

**TCR browser (**https://rstats.immgen.org/tcrbrowser/ )

The TCR browser displays the characteristics of the TCRs discovered by the immgenT project, their source and composition. It allows users to enter a particular TCR composition (V, J or CDR3 composition) to see if this or similar TCRs have been encountered in the immgenT dataset.

**immgenT Skyline (**https://rstats.immgen.org/Skyline/skyline.html?datagroup=immgenT%20Pseudobulk) The classic bar histograms in the ImmGen Skyline viewer represent the expression of a selected gene across all immgenT clusters, with weblinks to the gene’s characteristics and functions.

**Cluster characteristics (**https://www.immgen.org/ImmGenT/)

The immgenT Portal provides interactive visualization and information for level 2 clusters in the main lineages. Users can select individual level2 clusters to retrieve integrated biological information related to each cluster:

*1. Anatomical Distribution*: A mouse organ map visualizing the organs in which cells of this cluster are most represented.
*2. Sample Representation*: A list of the top 10 samples that most contributed to this cluster, providing context on the experimental conditions most associated with the cluster.
*3. Differentially expressed genes:* The volcano plots compare cells of the selected cluster to all other cells in the lineage (“vs. All) or to other cells in the same resting / experienced subgroups.

**Flow cytometry panels**

From the CITEseq data reporting on the expression of cell surface markers, immgenT groups have devised panels of markers for use in multi-parameter flow cytometry, and whose patterns can serve as proxies for the clusters in the immgenT framework, for CD4+, CD8+ and Treg lineages. The panels are listed, along with examples of stained populations.

#### Extended Data Note 3 - Cluster Robustness

To assess the robustness of immgenT clusters, we conducted several complementary analyses that leverage the multimodal nature of the dataset. Robust clusters are expected to exhibit reproducible transcriptional and protein expression profiles, as well as consistent distribution patterns across samples, organs, and experiments.

First, we examined cluster distribution across samples and experiments. No cluster was restricted to a single experiment (**Extended Data Fig. 4a**), indicating minimal batch effects. While some clusters were enriched in specific experiments, this reflected biological sampling differences rather than technical artifacts, such as enrichment of double-positive (DP) T cell clusters in thymus-containing experiments. Using the standardized spleen control spiked into each experiment, we further observed highly consistent cluster proportions across experimental replicates (**Extended Data Fig. 4b**), demonstrating both precision and reproducibility of the clustering framework.

Second, from a transcriptomic perspective, we quantified cluster distinctiveness within each dataset using a cluster-level separability score (Methods; **Fig. 4c**). For each dataset, this score compares the mean intra-cluster transcriptional distance to the mean distance between the cluster centroid and all other cluster centroids, providing a measure of global transcriptional distinctiveness that is robust to cluster size, local neighborhood structure, and the intrinsic continuity of T cell states. Scores are bounded between −1 and 1, with positive values indicating that cells within a cluster are, on average, more transcriptionally similar to each other than to cells in other clusters. Across 3,511 cluster–experiment combinations containing at least 10 cells, 99.5% exhibited positive separability scores, indicating that the vast majority of clusters represent reproducible and transcriptionally distinct cell states across datasets. Proliferative clusters showed the lowest separability, consistent with the ability of cells from multiple lineages to enter the cell cycle.

As a complementary analysis, we assessed the conservation of transcriptional programs across experiments by computing cosine similarity of gene expression profiles for each cluster across datasets (**Methods; Extended Data Fig. 4c**). All clusters except DP.wO (likely reflecting doublets) displayed high cross-experiment similarity, further supporting the reproducibility of these transcriptional states.

Cluster robustness was further evaluated in the protein expression space. Despite the CITEseq panel comprising only 128 surface proteins, RNA-defined clusters remained distinguishable in protein-based embeddings (illustrated in **Fig. 1d,e**). Building on this observation, we computed cluster-level separability scores in the protein PCA space for each dataset. Across 3,043 cluster–experiment combinations with at least 10 cells and available CITEseq data, 95% of clusters exhibited positive separability scores (**Extended Data Fig. 4d**), indicating the presence of conserved protein signatures across experiments. These protein-level signatures are described in greater detail in companion manuscripts.

Together, these analyses demonstrate that immgenT harmonizes experiment-specific clustering into a compact set of 107 consensus T cell states that are stable across experiments and robust across both RNA and protein modalities, providing a reliable framework for cross-study comparison and downstream biological interpretation.

#### Extended Data Note 4 – T-RBI reference-based integration

T-RBI (T cell Reference-Based Integration) was developed to make the immgenT molecular framework directly reusable for the analysis of external single-cell RNA-seq datasets. Conventional single-cell analyses generally generate a new dimensionality-reduction map and a new set of cluster annotations for every dataset, making direct comparisons across studies difficult. T-RBI instead maps query cells onto the fixed immgenT reference, providing a common coordinate system and annotation framework.

#### Integration and annotation

T-RBI uses transcriptomic data only and combines scVI/scANVI with anchored Minimum Distortion Embedding (MDE). Query cells are first filtered using a T-cell gene-expression score, and γδ T cells are identified separately from αβ T cells. Reference and query cells are then jointly modeled with scVI to generate a shared latent representation. scANVI is subsequently used to assign query cells first to major T-cell lineages (level 1) and then to the corresponding immgenT clusters (level 2). Each assignment is associated with a prediction confidence score. Cells with confidence scores <0.85 are initially designated “not classified”. A query-specific fine-tuning step then uses confidently annotated query cells to refine the assignment of these cells, while retaining cells that remain ambiguous as unclassified.

T-RBI therefore provides two complementary types of information for every query cell: an annotation within the immgenT hierarchy and a position within the immgenT reference maps. Query cells are projected onto the fixed all-T and lineage-specific MDEs using an anchored optimization in which the coordinates of the original immgenT cells remain unchanged. Thus, independent datasets can be displayed on exactly the same reference coordinates rather than on separately computed UMAPs. Importantly, positioning within the MDE and cluster annotation are distinct operations: a query cell can be positioned in the reference map without necessarily receiving a confident cluster assignment.

#### Detection of states not represented in the reference

A potential limitation of any reference-mapping approach is that a query cell might be forcibly assigned to the closest available reference population even when the corresponding state is absent from the reference. T-RBI addresses this in two ways. First, cells are left unclassified when their scANVI prediction confidence remains below the annotation threshold. Second, these unclassified cells are evaluated with a “discovery score” designed to identify coherent populations that are poorly represented by the reference. The discovery score is calculated independently of the immgenT integration, in the query dataset’s native PCA space. For each unclassified cell, the mean distance to its ten nearest annotated cells is compared with the mean distance to its ten nearest unclassified cells. Scores >1 indicate preferential proximity to other unclassified cells, and a conservative threshold of 1.1 is used to flag candidate states that may not be represented in the immgenT reference. Thus, isolated cells lying at boundaries between known states are distinguished from groups of unclassified cells that form a coherent transcriptional population.

We tested this behavior directly by mapping non-T cells (**Extended Data Figure 6b,g**), which clustered clearly away from the T-cell reference. We then examined more subtle T-cell states using cluster ablation (**Extended Data Fig. 7**). We removed individual CD4 clusters from the immgenT reference and mapped their cells back as query cells. Because MDE preserves local transcriptional neighborhoods, ablated cells were positioned near the transcriptionally closest remaining clusters. Thus, they generally returned to the region of the MDE previously occupied by the removed cluster, creating a population within the corresponding “hole” in the ablated reference (**Extended Data Fig. 7a**). Many ablated cells remained unclassified, whereas cells reassigned to neighboring clusters generally had reduced annotation confidence (**Extended Data Fig. 7b,c**). Because fine-tuning tended to increase reassignment to neighboring clusters, T-RBI reports annotations and confidence scores both before and after fine-tuning. The discovery score also increased after ablation, as expected when the corresponding reference cells were missing, with a substantial fraction of cells exceeding the 1.1 threshold (**Extended Data Fig. 7d**). Thus, T-RBI does not simply force cells into the nearest available cluster when their original state is absent. This analysis also provides an independent test of the immgenT clustering itself: if two neighboring clusters represented arbitrary subdivisions of the same state, removing one should result in high-confidence reassignment of its cells to the other. The incomplete and lower-confidence reassignment instead supports the molecular distinctiveness of these clusters.

#### Validation with external datasets

We evaluated T-RBI on 16 independent studies comprising 335,042 cells (**Fig. 5, Extended Data Fig. 6, Extended Data Table 5**). These studies included conventional and unconventional T-cell populations, previous T-cell atlases, CAR T cells, Tregs from tissues not represented or sparsely represented in immgenT, and diverse infectious and inflammatory settings. The published datasets mostly used 3′ and 5′ 10X Genomics chemistries, but we also tested newer single-cell protocols from unpublished data, including Parse, 10x Flex, and 10x single-nucleus RNA-seq, which also mapped correctly to the reference. More than 99% of bona fide T cells received confident lineage-level annotations, with positioning concordant with expression of canonical lineage markers such as *Cd4, Cd8a, Cd8b1, Foxp3, Zbtb16,* and *Trdc*. At cluster resolution, a mean of 97% of T cells (range 85–99% across datasets) were confidently annotated. Unclassified T cells were generally interspersed among annotated cells rather than forming separate populations, and consequently showed low discovery scores. In contrast, contaminating non-T cells provided a positive control: they remained spatially separated from the immgenT T-cell manifold and exhibited high discovery scores.

#### T-RBI mapping preserves information present in the original datasets while placing it into a common framework

For example, the five populations originally identified in the Miller et al. chronic LCMV dataset remained spatially distinct after mapping, while immgenT annotations further resolved them into multiple CD8 states (**Fig. 5a–c**). Thus, reference mapping does not require erasing the original annotation or organization; rather, the original and immgenT annotations can be compared directly on the same cells.

#### T-RBI provides an alternative to defining T-cell states from individual markers or fixed gene signatures

Marker combinations and signatures can be highly context-dependent, whereas T-RBI uses high-dimensional relationships across the transcriptome, including nonlinear combinations of genes. This is illustrated by tissue-resident memory CD8 T cells (**Fig. 10**): canonical CD69/CD103 expression and a published Trm gene signature were not specific for the molecular state occupied by bona fide LCMV memory cells, whereas T-RBI mapped the cells used to derive the published Trm signature predominantly to CD8.Q. The same CD8.Q state contained memory cells generated in several tissues and immune challenges, illustrating how reference-based annotation can recognize a shared molecular state across contexts in which individual markers may differ.

#### Public implementation and outputs

T-RBI is publicly available through the immgenT portal (https://rstats.immgen.org/immgenT_integration_analysis/) and can map datasets against either the complete T-cell reference or selected lineages. Users provide a raw count matrix, feature and barcode information, and sample metadata containing batch information; known lineage information can also be supplied when available. The current implementation has been tested primarily with 10X Genomics 3′ and 5′ scRNA-seq datasets and has been successfully applied to other formats, including Flex and Parse datasets. To manage computational resources, the web implementation currently accepts up to 25,000 cells per submission; larger datasets can be divided into multiple submissions.

After processing, T-RBI returns a PDF summary and cell-level output files containing the original cell identifiers, level 1 and level 2 immgenT annotations, annotation confidence scores before and after fine-tuning, discovery scores for unclassified cells, and coordinates within the all-T and lineage-specific MDEs. The report also displays confidence-score distributions, unclassified cells in a dataset-specific UMAP, and discovery scores within the all-T MDE. These outputs allow the user both to adopt the immgenT annotations and to inspect cells for which the reference assignment is uncertain.

Together, T-RBI turns immgenT from a static atlas into a reusable reference system: external datasets can be positioned and annotated within a fixed molecular coordinate system while retaining the ability to identify states not adequately represented in the current reference. With user permission, submitted datasets may also contribute to future extensions of the reference, allowing the immgenT framework to evolve as additional T-cell states and experimental contexts are sampled, in the open-source tradition of this project.

**Extended Data Figure 1. Quality control metrics.**

**(a)** Box plots showing, for each experiment (IGT), the percentage of cells retained after quality control and the percentage of cells removed due to low gene counts (RNA nGenes), high mitochondrial RNA content (RNA dead cells), low protein counts, high non-specific protein signal, or non-T-cell identity, relative to the pre-QC cell number.

**(b)** Number of cells per experiment (IGT) and per sample, identified by unique hashtag (HT) within each experiment. Bars are colored by HT.

**Extended Data Figure 2. CITEseq quality control (128-plex).**

**(a–c)** RNA–protein correlations for representative surface markers with high dynamic range and performance (CD4, CD8β, Ly49A) **(a),** intermediate dynamic range (CD127) **(b),** and poor performance due to lack of detectable protein signal (CCR7, FASL) **(c)**. Points are colored by T-cell lineage.

**(d)** Summary of all antibodies in the 128-plex CITEseq panel, plotted according to protein dynamic range (log10 protein max minus protein min, x-axis) and Spearman correlation between RNA and protein expression (y-axis)

**Extended Data Figure 3. Spleen standards across experiments; T-cell lineage separation.**

**(a)** All-T MDE highlighting spleen standard samples, colored by experiment (IGT), demonstrating overlap of standards across experiments.

**(b)** All-T MDE highlighting each major T-cell lineage separately.

**(c)** Expression of TCRγδ versus TCRβ, CD4 versus CD8β, and CD8β versus CD8α (CITEseq, log1p(CP10K)) shown separately for each T-cell lineage.

**(d)** all-T MDE with iNKT cells identified by their canonical invariant TCR, defined by TRAV11–TRAJ18 usage and characteristic CDR3α junction sequences (CVVGDRGSALGRLHF, CVVADRGSALGRLHF, CVVVDRGSALGRLHF).

**(e)** all-T MDE with MAIT cells identified by their canonical TCR, defined by TRAV1–TRAJ33 usage with a characteristic CDR3α length of 12 amino acids.

**(f)** All-T MDE highlighting proliferating cells (black) and miniverse (wM) cells (red).

**Extended Data Figure 4. Cluster robustness metrics**

**(a)** Stacked bar plot showing the experiment (IGT) composition of each immgenT consensus cluster, demonstrating that no cluster is experiment-specific (except DP clusters in thymus experiments).

**(b)** Box plots showing the distribution of immgenT clusters across spleen standard samples, illustrating reproducibility across experiments.

**(c)** Distribution of cosine similarity scores for gene expression profiles of cells assigned to the same cluster across datasets.

**(d)** Distribution of cluster separability scores for each immgenT consensus cluster in protein expression space. Scores > 0 indicate proteomically distinct clusters. Colors indicate experiment (IGT).

**Extended Data Figure 5. Distribution of stress-related transcriptional signatures across the immgenT atlas.**

**(a–h)** All-T MDE showing single-cell scores for representative signatures: **(a)** 1-h dissociation and **(b)** 2-h dissociation signatures from van den Brink et al.^39^; **(c)** immediate-early response signature from Bahrami et al.^108^; **(d)** integrated stress response signature from Alicea Pauneto et al.^110^; **(e)** ER-stress signature (GO:0034976); **(f)** interferon-stimulated gene signature from ImmGen^109^; **(g)** hypoxia signature (GO:0001666); and **(h)** mitochondrial genes. Higher scores are indicated by increasing color intensity.

**(i)** Heatmap showing mean scores for an expanded set of stress-related signatures across immunological conditions, including the ER-stress signature from Chow et al.^111^, the hypoxia signature from Bono et al.^112^, and the immediate-early response (IER) signature from Bahrami et al.^108^ (*Nr4a1, Egr1, Egr2, Jun, Junb, Fos,* and *Fosb*). Values are shown as mean-centered scores.

**(j)** Heatmap showing mean signature scores across anatomical sites. Values are shown as mean-centered scores.

**(k)** Heatmap showing mean signature scores across immgenT clusters, grouped by T-cell lineage. Interferon-related signatures show selective enrichment in specific states, including CD8.F and CD4.D, whereas most other stress-related programs are distributed more broadly across clusters, conditions, and tissues. Values are shown as mean-centered scores.

**Extended Data Figure 6. T-RBI metrics**

**(a)** UMAP of cells from Miller *et al.*^64^ showing the immgenT unclassified cells (red) intermingled with annotated cells (grey).

**(b)** All-T MDE with T cells from 16 external studies (**Extended Data Table 5**) integrated by T-RBI. immgenT cells shown in the background (grey). Non-T cells from the external studies (red) cluster separately, demonstrating that integration does not force overlap with T cells.

**(c)** Bar plot showing the proportion of cells from external studies that are not annotated with a T-cell lineage (level-1 annotation).

**(d)** All-T MDE showing expression of *Cd4, Cd8b1, Cd8a, Trdc, Foxp3,* and *Zbtb16* in the external studies, illustrating concordance between lineage marker expression and immgenT positioning.

**(e)** Bar plot showing the proportion of T cells from external studies that are not annotated with an immgenT cluster (level-2 annotation).

**(f)** All-T MDE with integrated external T cells (foreground) and immgenT cells (grey background). Unannotated T cells at the cluster level (red) remain embedded within the immgenT manifold and do not form distinct clusters.

**(g)** All-T MDE colored by the single-cell discovery score for external studies. Non-T cells exhibit high discovery scores (red; > 1.1).

**(h)** Distribution of discovery scores across external studies, including non-T cells as positive controls. High scores indicate query cells that form coherent states outside the immgenT reference; non-T cells show consistently elevated discovery scores.

**Extended Data Figure 7. Ablation tests assess the ability of T-RBI to detect T-cell states absent from the reference.**

To simulate mapping of novel T-cell states, individual CD4 reference clusters were iteratively removed from the immgenT reference, and cells belonging to the omitted cluster were mapped back as query cells using T-RBI. Results were compared with mapping against the complete, non-ablated reference.

**(a)** CD4 MDEs showing the positions of query cells from each omitted cluster after T-RBI against the ablated reference. Query cells map predominantly to the region occupied by the corresponding cluster in the complete reference, thereby filling the “hole” created by its removal.

**(b)** T-RBI cluster assignments of cells from each omitted cluster. Rows indicate the original cluster and columns the assigned cluster after ablation; dot size and color indicate the proportion of cells receiving each assignment. Values in the ‘not classified’ column indicate the percentage of cells remaining unannotated.

**(c)** T-RBI annotation confidence scores for cells misclassified after ablation, shown before and after annotation fine-tuning and compared with scores for the same cells mapped to the complete reference without ablation.

**(d)** Single-cell discovery scores for cells from each omitted cluster with the complete reference (black) or after ablation (red). The dashed line indicates the discovery-score threshold of 1.1 used to flag candidate states not represented in the reference.

**Extended Data Figure 8. Cluster distributions across tissues.**

All-T MDE showing the distribution of T cells at baseline in bone marrow, peritoneal cavity, submandibular gland, uterus, and placenta, colored by lineage.

**Extended Data Figure 9. Effector molecule expression in immgenT clusters**

**(a)** Heatmap showing log1p(CP10K) expression of the 100 most cluster-specific effector genes, defined as genes reaching at least 0.5 log1p(CP10K) expression in at least one immgenT cluster. Rows (genes) and columns (clusters) are ordered by hierarchical clustering. Annotation bars indicate T-cell lineage and activation status (resting versus activated).

**(b)** Bar plot showing average expression of Il9 across immgenT clusters.

**Extended Data Figure 10. Transcription factor expression in immgenT clusters**

**(a,b)** Heatmaps showing log1p(CP10K) expression of the 20 most specific transcription factors across T-cell lineages (a) and across immgenT clusters (b).

**(c–f)** Bar plots showing average expression of *Sox4* **(c),** *Zbtb32* **(d),** *Tshz2* **(e),** and *Hlf* **(f).** across immgenT clusters

**Extended Data Figure 11. Markers and gene signatures across immgenT states**

**(a)** CD4-specific MDE highlighting Th1 (CD4.R/S), Th2 (CD4.T/U), and Th17 (CD4.X/Y) “tip” clusters.

**(b)** CD4-specific MDE showing expression of *Cxcr3, Il12rb2, Ccr4, Il4ra, Ccr6,* and *Il23r*

**(c)** All-T MDE showing expression of the core Treg gene signature^90^ across all immgenT T cells.

**Extended Data Table 1. Sample-level metadata for the immgenT dataset**

**Extended Data Table 2. CITEseq panel information**

**Extended Data Table 3. Quality control metrics for each sample**

**Extended Data Table 4. Quality control metrics for the CITEseq panel**

**Extended Data Table 5. List of integrated studies by T-RBI**

**Extended Data Table 6. Transcription factor specificity score (AUC) in T cell lineages**

**Extended Data Table 7. Transcription factor specificity score (AUC) in T cell clusters**

**Extended Data Table 8. Correspondence between immgenT clusters and common annotations**

