## Extended Data Figures 1-11 for "immgenT: A Comprehensive Reference of Convergent T-cell States in the Mouse"

Extended Data Figure 1

a

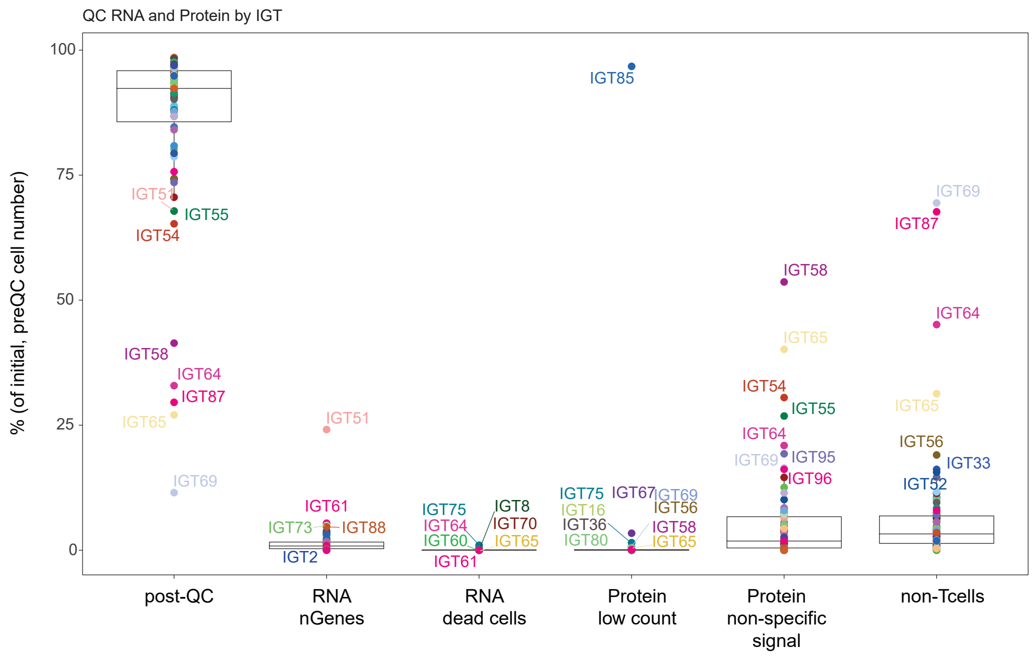


b

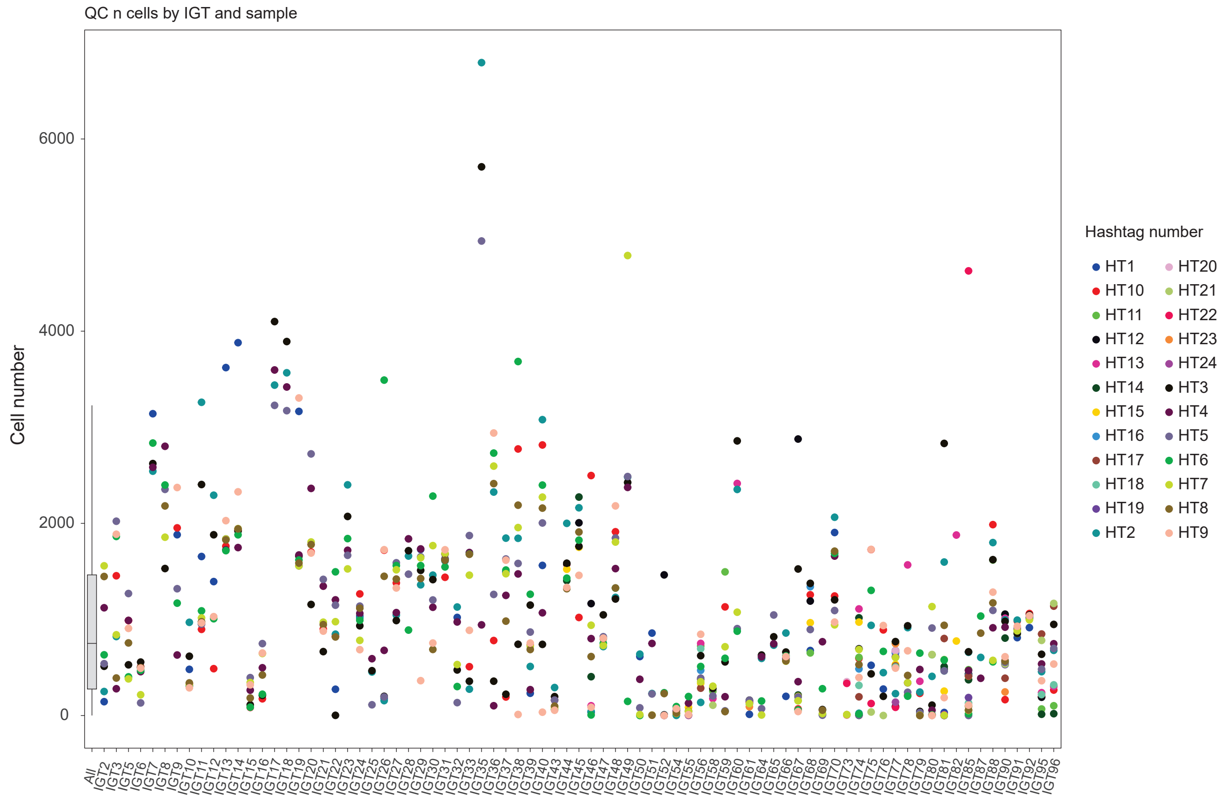

Extended Data Figure 2

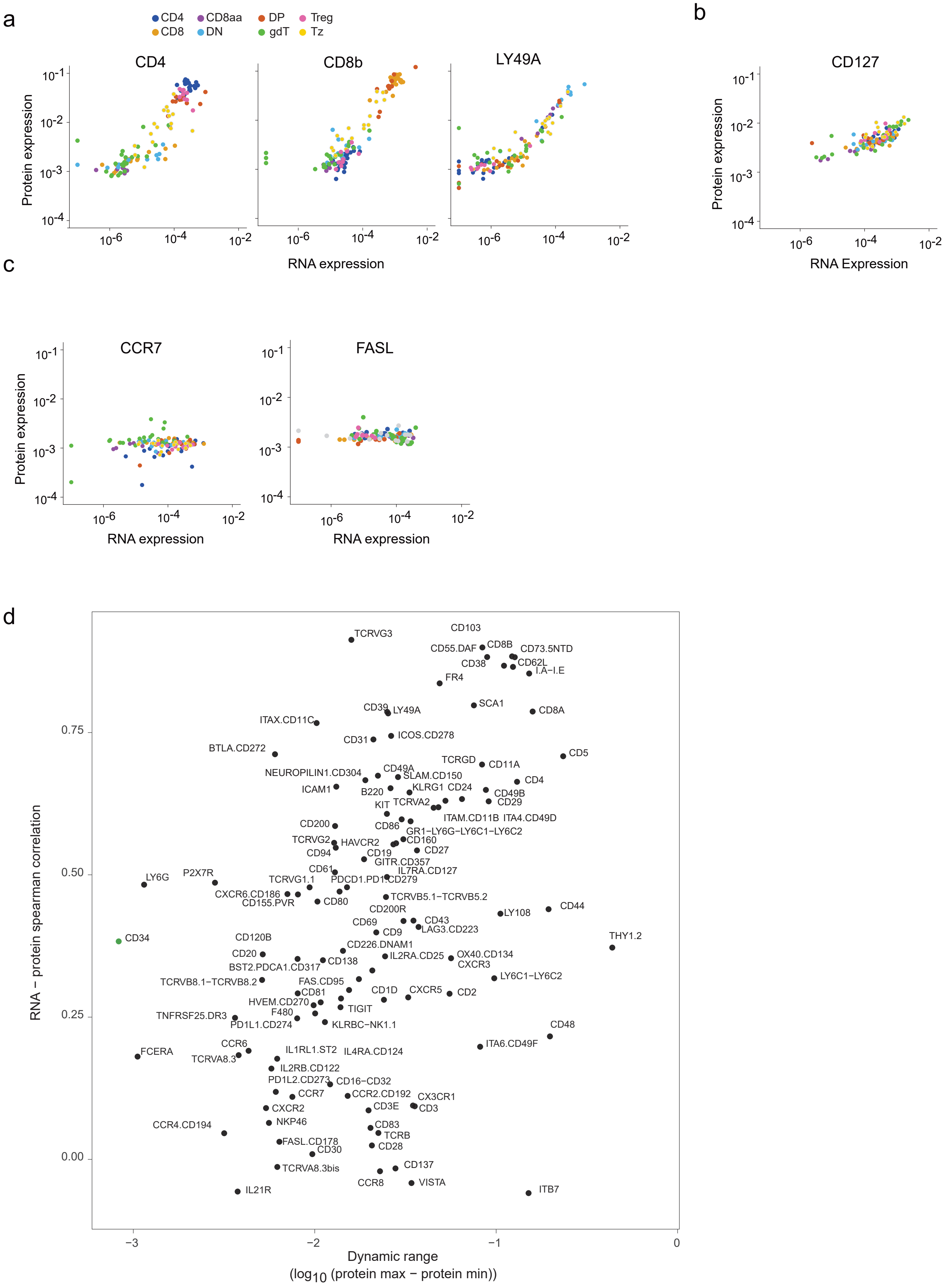

Extended Data Figure 3

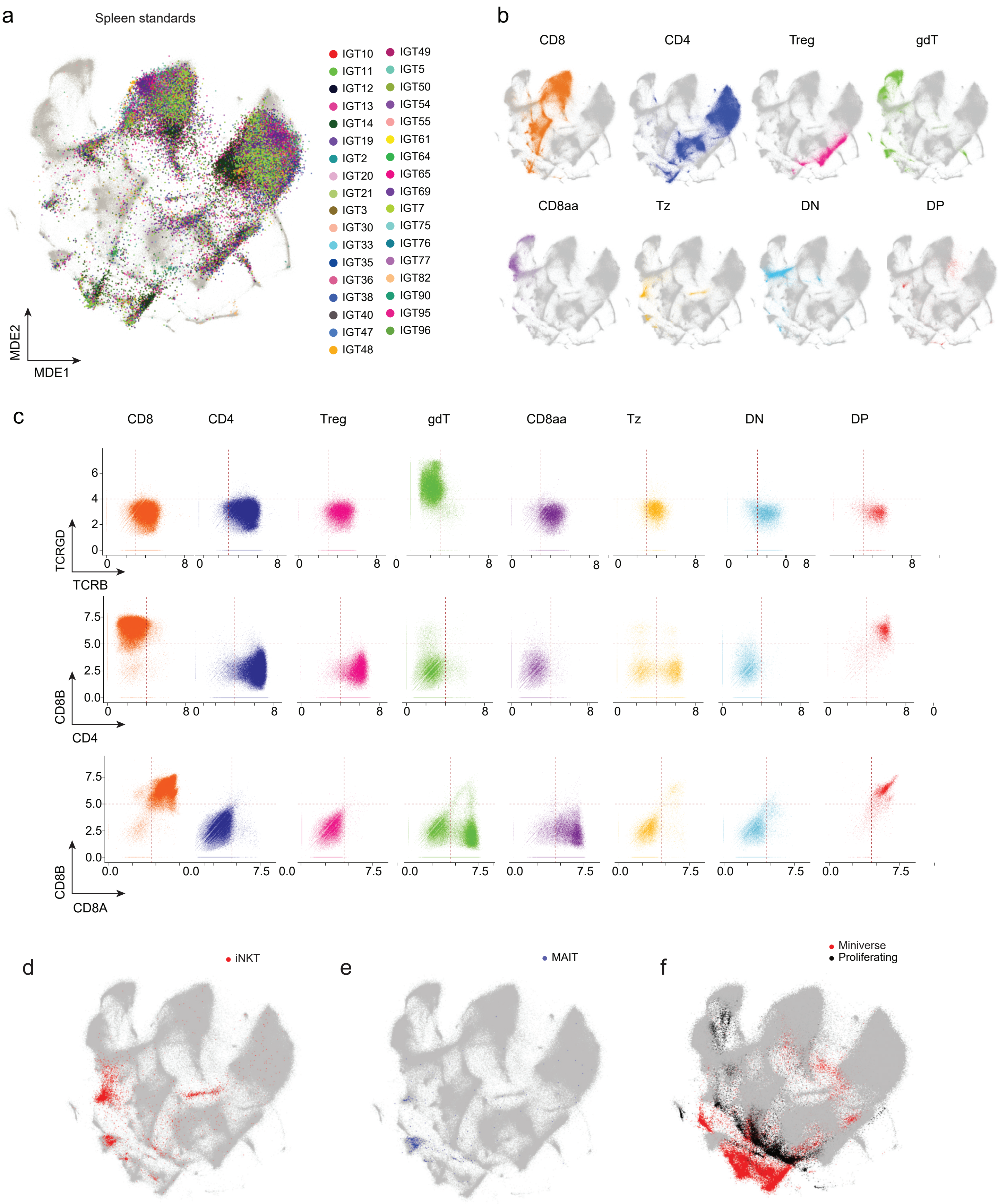

Extended Data Figure 4

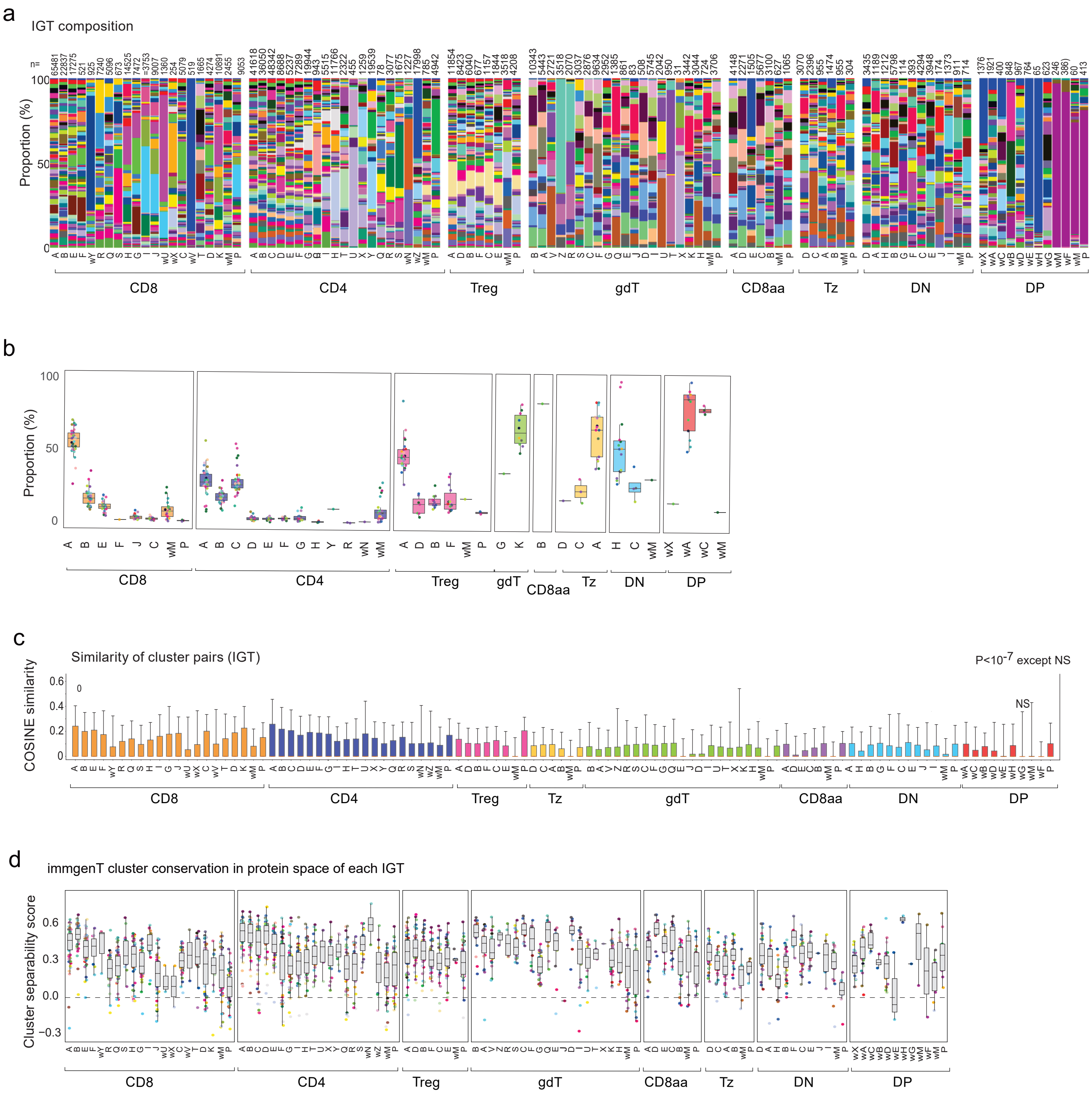

### Extended Data Figure 5

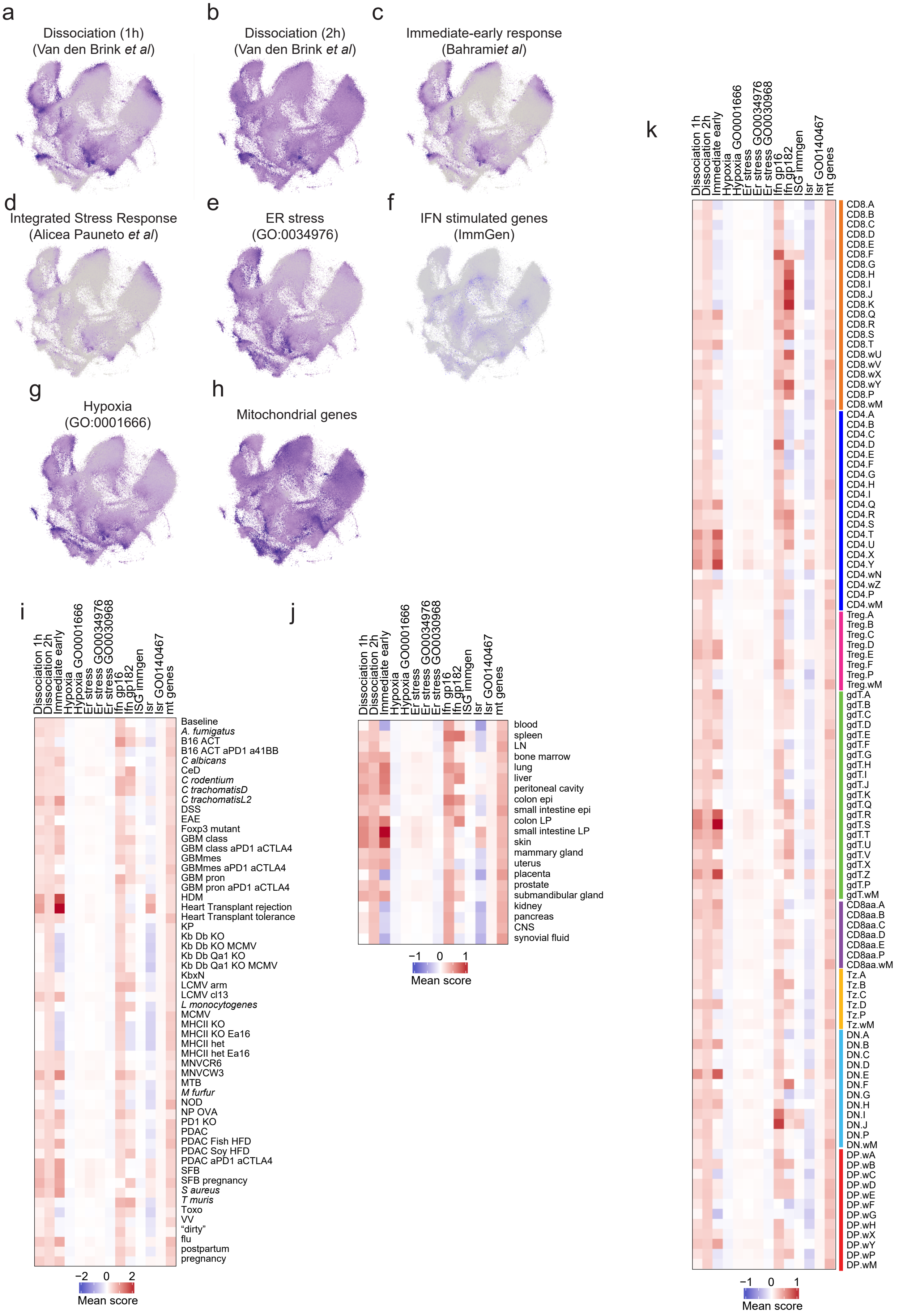

Extended Data Figure 6

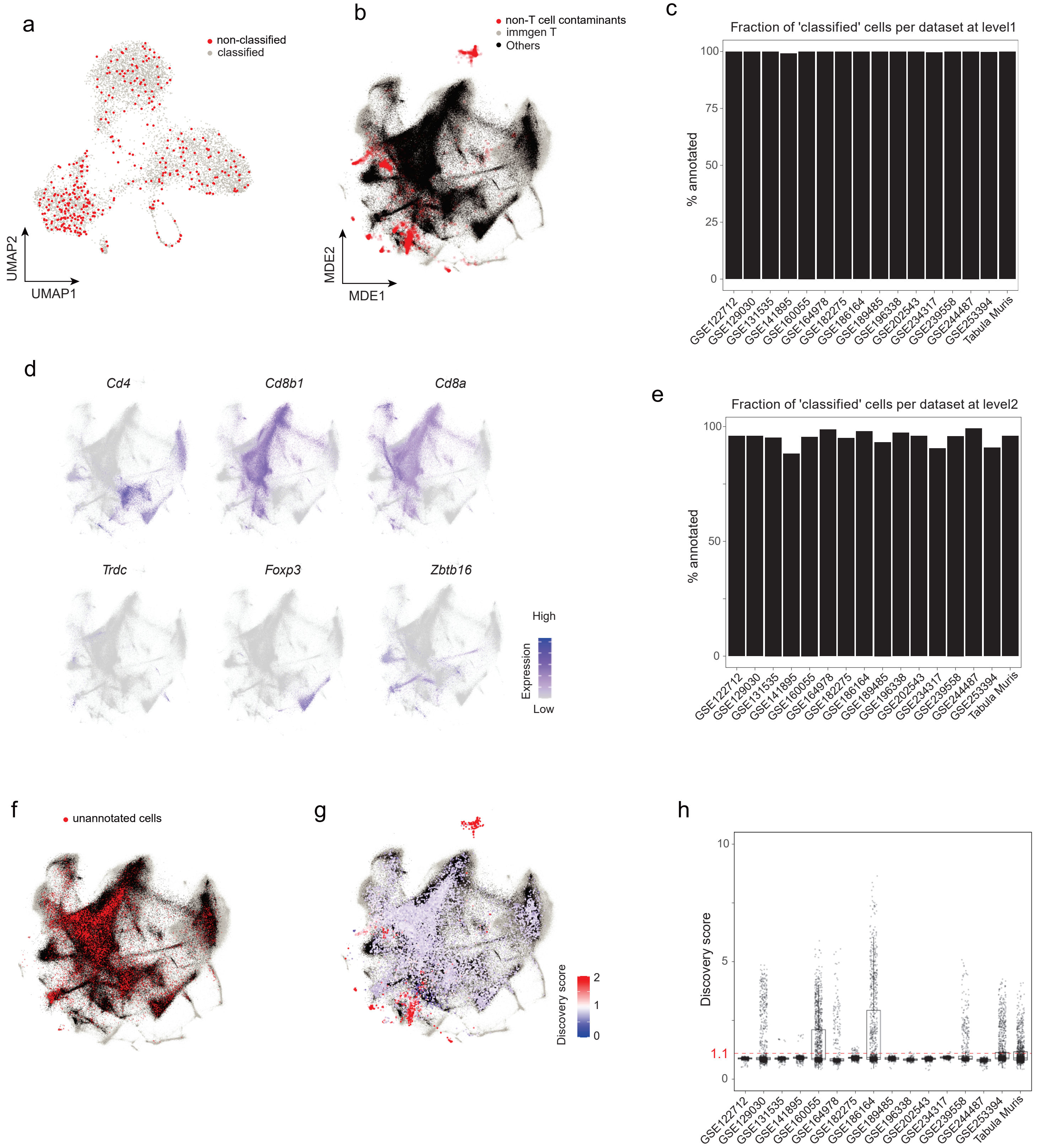

#### Extended Data Figure 7

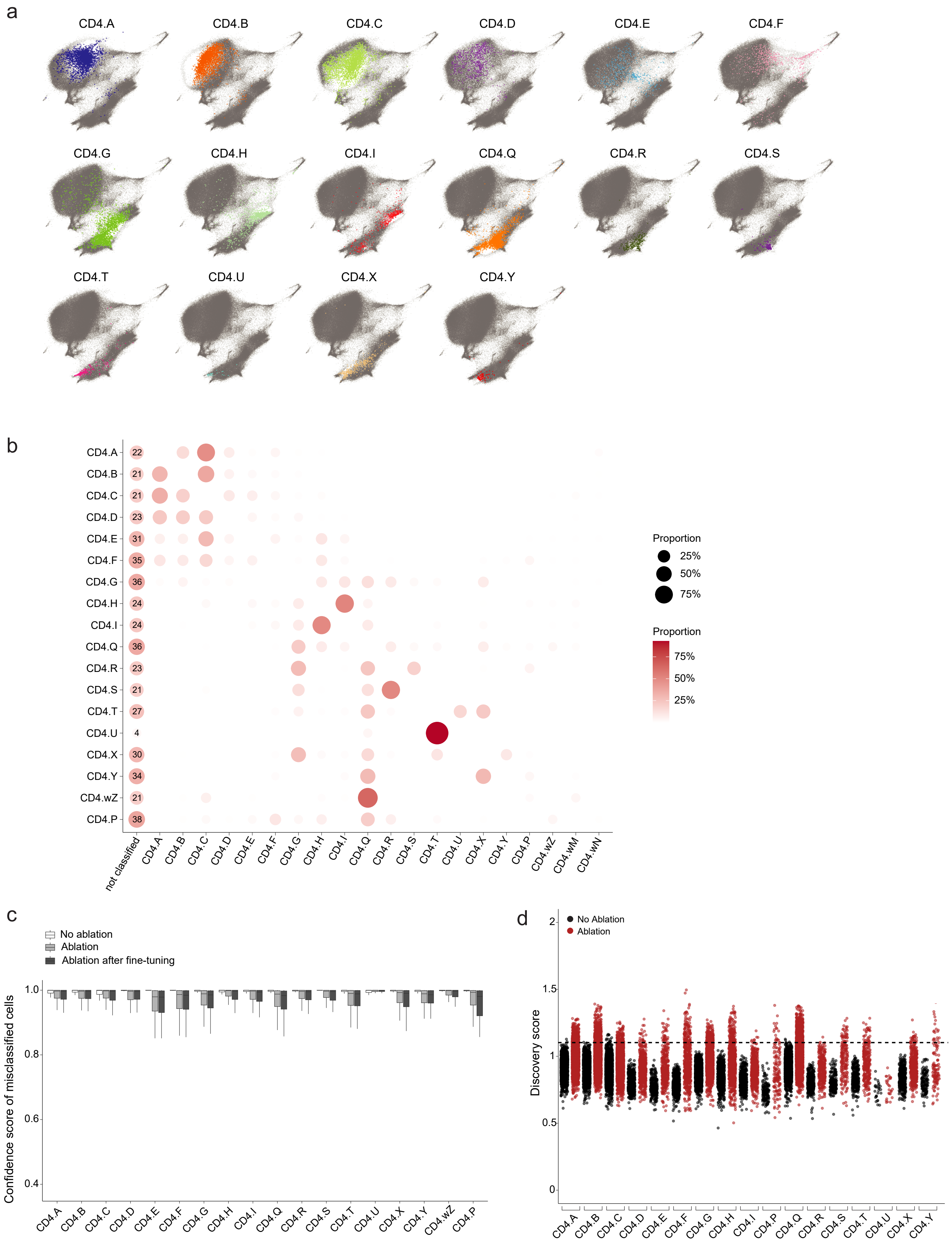

### Extended Data Figure 8

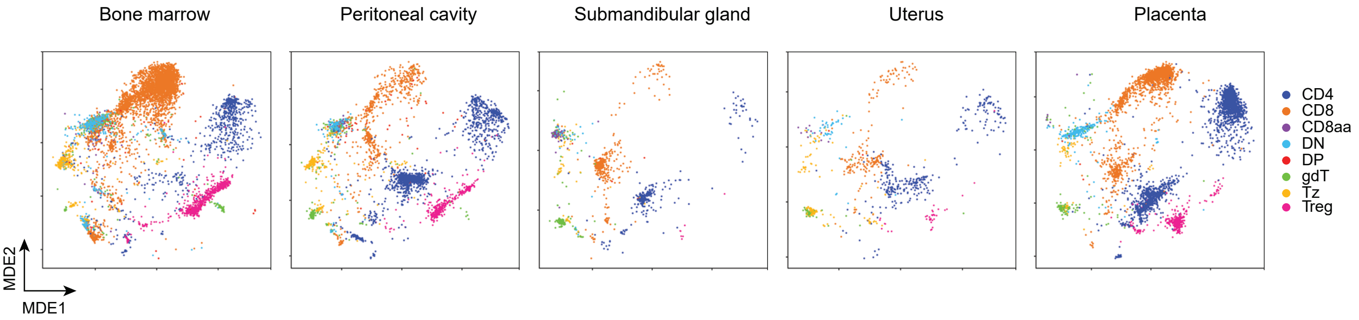

### Extended Data Figure 9

a

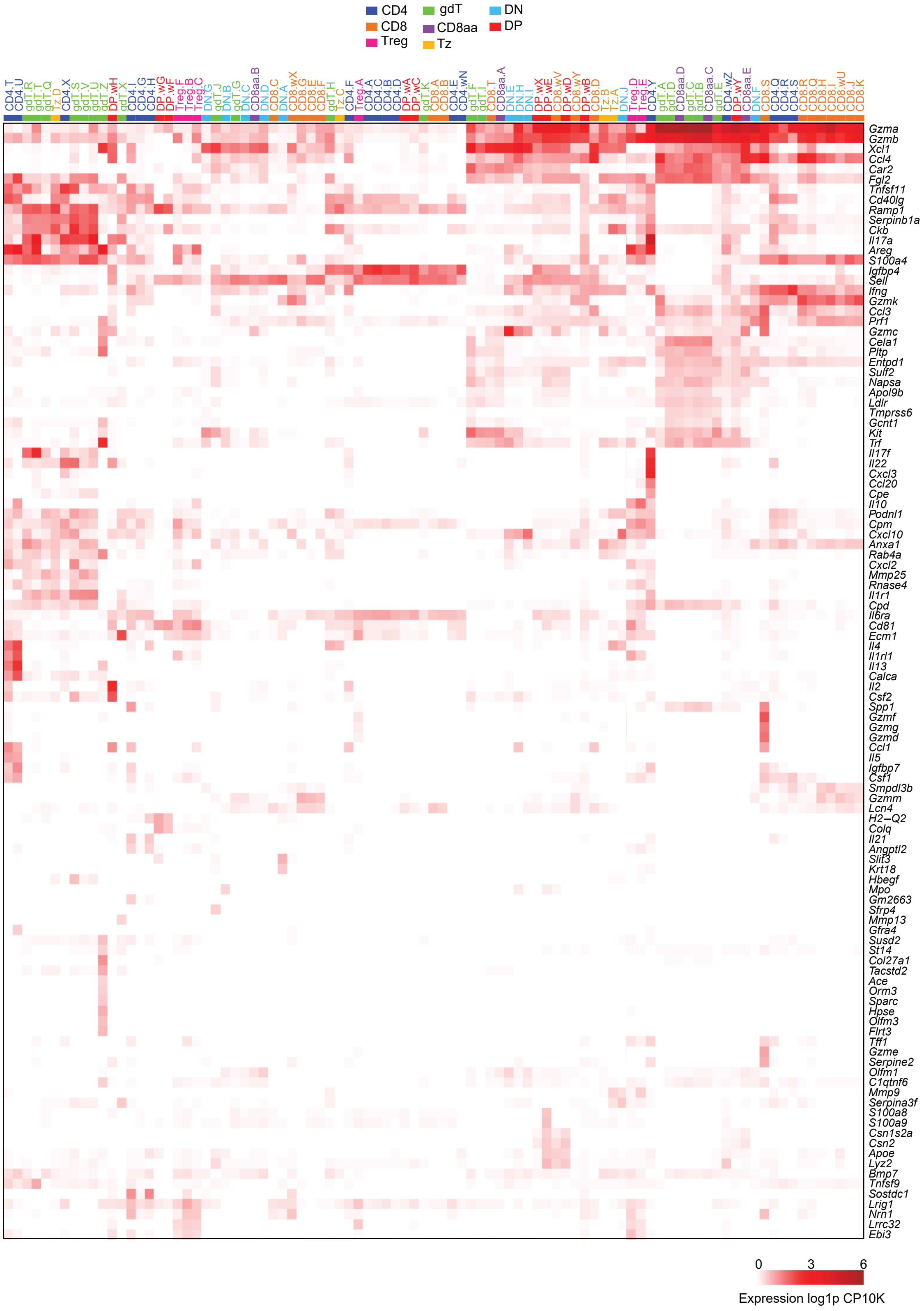


b

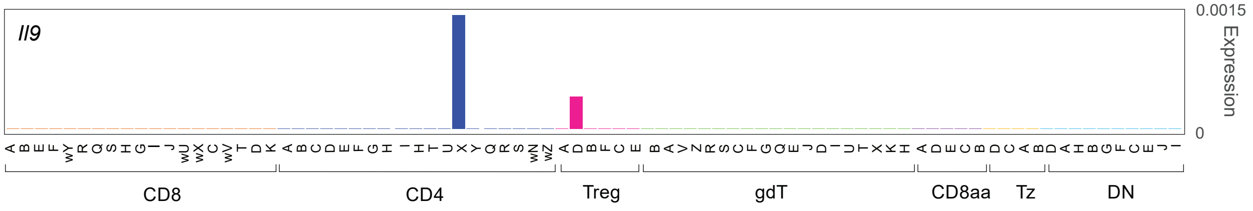

Extended Data Figure 10

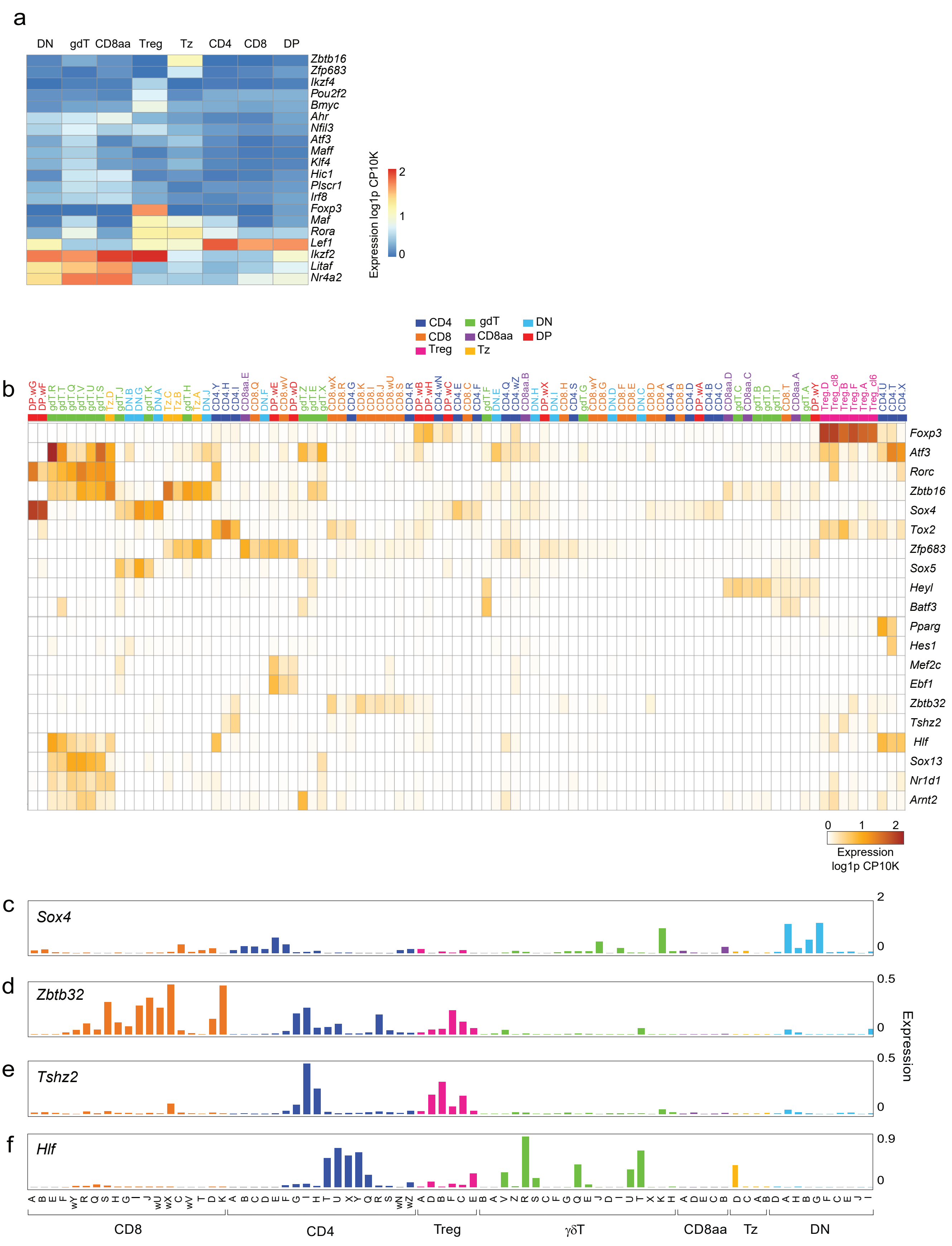

Extended Data Figure 11

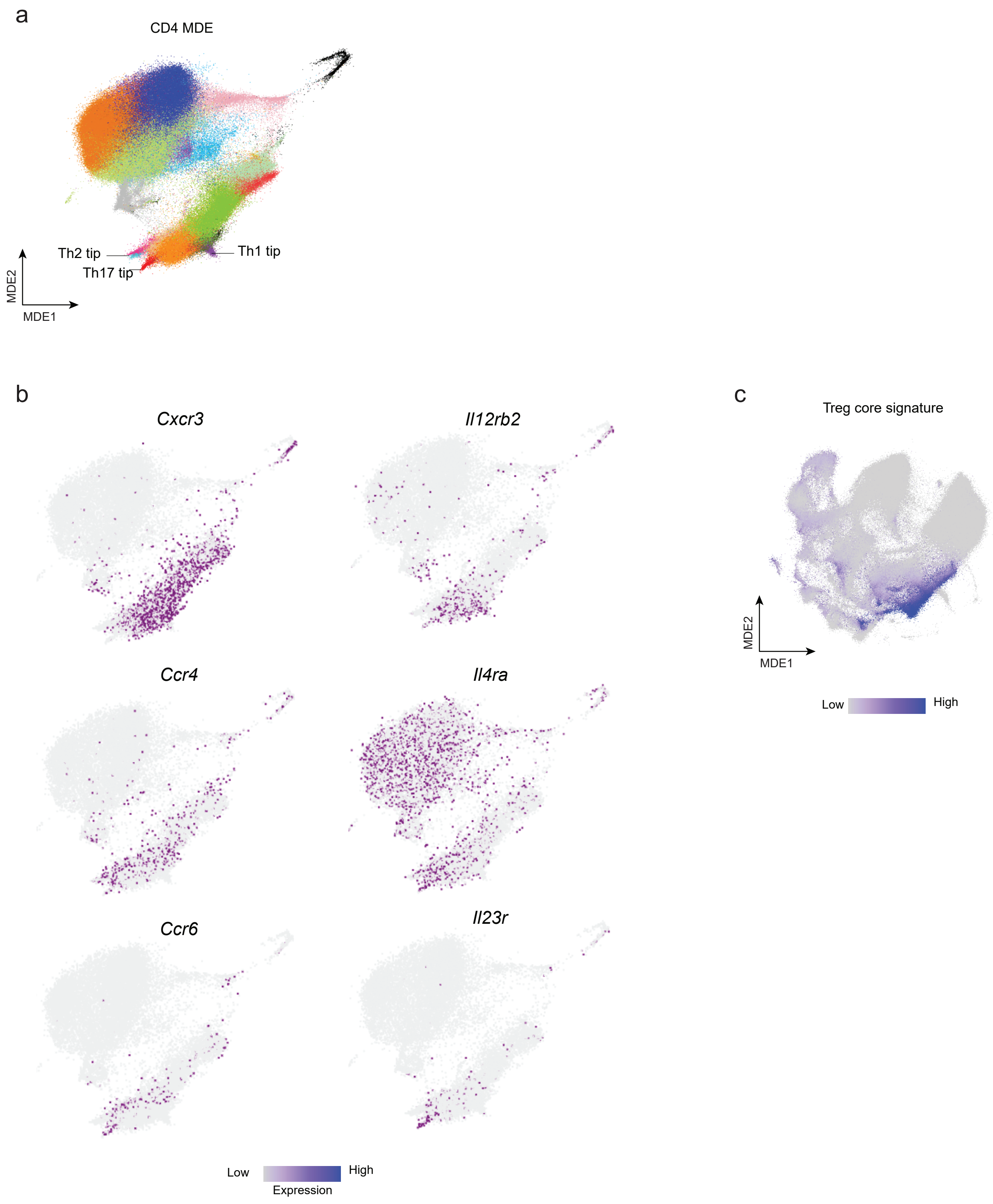
